# Impaired α-tubulin re-tyrosination leads to synaptic dysfunction and is a feature of Alzheimer’s disease

**DOI:** 10.1101/2021.05.17.443847

**Authors:** Leticia Peris, Xiaoyi Qu, Jean-Marc Soleilhac, Julie Parato, Fabien Lanté, Atul Kumar, Maria Elena Pero, José Martínez-Hernández, Charlotte Corrao, Giulia Falivelli, Floriane Payet, Sylvie Gory-Fauré, Christophe Bosc, Marian Blanca Ramírez, Andrew Sproul, Jacques Brocard, Benjamin Di Cara, Philippe Delagrange, Alain Buisson, Yves Goldberg, Marie-Jo Moutin, Francesca Bartolini, Annie Andrieux

**Author notes:** Equal contribution (Leticia Peris, Xiaoyi Qu). These authors contribute equally to the study (Marie-Jo Moutin, Francesca Bartolini, Annie Andrieux). Corresponding authors : Leticia Peris, Marie-Jo Moutin, Francesca Bartolini and Annie Andrieux.

## Abstract

In neurons, dynamic microtubules play regulatory roles in neurotransmission and synaptic plasticity. While stable microtubules contain detyrosinated tubulin, dynamic microtubules are composed of tyrosinated tubulin, suggesting that the tubulin tyrosination/detyrosination (Tyr/deTyr) cycle modulates microtubule dynamics and synaptic function. In the Tyr/deTyr cycle, the C-terminal tyrosine of α-tubulin is re-added by tubulin-tyrosine-ligase (TTL). Here we show that TTL^+/−^ mice exhibit decreased tyrosinated microtubules, synaptic plasticity and memory deficits, and that reduced TTL expression is a feature of sporadic and familial Alzheimer’s disease (AD), with human APPV717I neurons having less dynamic microtubules. We find that spines visited by dynamic microtubules are more resistant to Amyloidβ_1-42_ and that TTL, by promoting microtubule entry into spines, prevents Aβ_1-42_-induced spine pruning. Our results demonstrate that the Tyr/deTyr cycle regulates synaptic plasticity, is protective against spine injury, and that tubulin re-tyrosination is lost in AD, providing evidence that a defective Tyr/deTyr cycle may contribute to neurodegeneration.

## INTRODUCTION

The neuronal microtubule (MT) cytoskeleton plays a fundamental role in the development and long-term maintenance of axons and dendrites. Research over the past two decades has revealed that dynamic MTs, in particular, critically contribute to synaptic structure and function within both pre- and postsynaptic compartments ^1, 2^. Dynamic MTs regulate synaptic vesicle cycling by providing paths for bidirectional transport between presynaptic terminals, a rate-limiting step in exocytosis at sites of release ^3–6^. In dendritic spines, while the core cytoskeletal structure consists of actin filaments, dynamic MTs originating in the dendritic shaft sporadically enter the spine head and directly impinge on the regulation of spine composition and morphology ^7, 8^. MT entry into spines is dependent on synaptic activity, Ca^2+^ influx, actin polymerization, and correlates with changes in synaptic strength ^9^. In cultured rodent hippocampal neurons and organotypic slices, stimulation of postsynaptic N-methyl-D-aspartate (NMDA) receptors by chemical long-term potentiation (LTP) protocols or glutamate photo-release leads to higher frequency and longer duration of spine invasions by MTs, concurrent with spine enlargement ^10–12^. Conversely, chemical induction of long-term depression (LTD) decreases MT invasions, indicating that MT targeting into spines is sensitive to plasticity signals ^13–15^. Spine invasions, as well as synaptic plasticity, specifically involve dynamic MTs, as both invasions and LTP are blocked when MT dynamics are inhibited by low doses of nocodazole ^11^ or taxol ^8, 16^. Consistent with these results, efficient contextual fear conditioning in mice appears to require transient accumulation of dynamic MTs at dentate gyrus synapses ^17^. Together, these findings indicate that changes in synaptic MT dynamics may affect both pre- and postsynaptic functions.

MT dynamics rely on the intrinsic capacity of MTs to alternate phases of polymerization and depolymerization. Various cellular factors have been shown to modulate MT dynamics including the nature of tubulin isoforms, GTP hydrolysis, MT associated proteins and also various post translational modifications (PTMs) of tubulin ^18, 19^. One prominent modification is the reversible removal of the C-terminal tyrosine residue of α-tubulin subunits, which is exposed at the external surface of MTs. This residue is cleaved off by specific tubulin carboxypeptidases (TCPs), such as the recently identified Vasohibin 1 (VASH1) - Small Vasohibin Binding Protein (SVBP) and Vasohibin 2 (VASH2)-SVBP complexes ^20–22^. When detyrosinated MTs depolymerize, the tyrosine is rapidly restored on disassembled α-tubulin by the enzyme tubulin tyrosine ligase (TTL), thereby replenishing the soluble tubulin pool with full-length subunits that are then available for renewed polymerization ^23–27^. Due to these sequential reactions, tubulin undergoes a continuous cycle of detyrosination and re-tyrosination (deTyr/Tyr). Detyrosinated MTs can be further processed by cytosolic carboxypeptidases of the deglutamylase family to generate Δ2 and Δ3 tubulins through the sequential cleavage of the final 2 or 3 amino acids, respectively ^28–31^. Δ2 tubulin cannot be re- tyrosinated by TTL, and is thus removed from the Tyr/deTyr cycle ^27, 32^. It follows that TTL suppression induces an accumulation of detyrosinated and Δ2 -tubulins, whereas TCP inhibition has the opposite effect ^33–35^.

Newly formed tyrosinated MTs are highly dynamic, contrary to detyrosinated MTs which are typically more stable ^26, 36, 37^. Indeed, while it is known that tubulin detyrosination can occur on previously stabilized MTs ^38, 39^, there is also evidence that detyrosination of tubulin may itself promote MT stability by protecting MTs from the depolymerizing activity of kinesin-13 motors ^40^. Thus, MT dynamics and the Tyr/deTyr cycle are intertwined, and modulation of the cycle is critical to processes in which MTs need to maintain a specific dynamic state. Moreover, MT detyrosination confers preferential binding for specific motors and other MT-associated proteins (MAPs), allowing tyrosination-dependent loading of selected cargoes and MT modulators. For example, in neurons, detyrosinated MTs play a unique role in neuronal transport by acting as preferential tracks for kinesin-1 and kinesin-2 ^41–51^ while inhibiting cytoplasmic linker proteins (CLIPs) and dynein loading onto MT plus ends ^33, 52^. Additional roles for detyrosinated MTs as regulators of MT severing enzymes have been suggested ^53, 54^. In neurons, these functions regulate the trafficking of cargos, axon outgrowth and branching. For example, Kinesis related protein 5 (KIF5)/kinesin-1 preferentially moves along detyrosinated MTs, and detyrosination affects its velocity in vitro ^51^. Kinesin-1 is involved in mitochondria trafficking ^55^, targeting of α-amino-3-hydroxy-5-methyl-4-isoxazolepropionic acid (AMPA) receptors to dendrites ^56^ and AMPA receptor-mediated synaptic transmission ^57^. KIF5/kinesin-1 may also regulate inhibitory transmission by directing the transport of Gamma aminobutyric acid (GABA) receptors via huntingtin-associated protein 1 (HAP1) ^58, 59^. Furthermore, robust kinesin-2 motility requires detyrosination of α-tubulin ^50^ and homodimeric KIF17/kinesin-2 has been implicated in the transport of GluN2B whereas disruption of KIF17/Kinesin-2 impaired LTP, LTD and cAM response element-binding protein (CREB) responses in mice ^60^.

While the function of Δ2 tubulin remains unknown, it is very abundant in neurons where it accumulates on very long-lived MTs ^32^. Unbuffered accumulation of Δ2 tubulin, however, has been recently associated with axonal degeneration that occurs following inhibition of mitochondrial motility ^35^. In the brain, significant alteration of the Tyr/deTyr cycle from birth modifies the relative ratio of tyrosinated, detyrosinated and Δ2 tubulin, leading to severe neurodevelopmental phenotypes in mice ^34, 61^. The *Svbp* knock-out (KO) in mice, which lead to no activity of the tubulin carboxypeptidases VASH1 and VASH2, resulted in a perturbed neuronal migration in the developing neocortex, microcephaly and cognitive defects, including mild hyperactivity, lower anxiety and impaired social behavior ^34^. Similarly, biallelic inactivating SVBP variants in human cause a syndrome involving brain anomalies with microcephaly, intellectual disability and delayed gross motor and speech development ^34, 62^. On the other hand, *Ttl* KO mice show disorganization of neocortical layers, disruption of the cortico-thalamic loop, and death just after birth ^61, 63^. However, it remains unknown whether post-developmental alteration of the Tyr/deTyr tubulin cycle plays a role in neurodegenerative disease.

Alzheimer’s disease (AD) is an age-related, neurodegenerative disorder, defining pathological features of which are overabundance of amyloid beta peptide (Aβ) and hyperphosphorylated tau ^64^. The most prominent clinical symptom is progressive memory loss, and decreases in synaptic density are associated with cognitive impairment ^65, 66^. AD is a multifactorial disease, with both genetic and environmental etiologies ^67^. The London (V717I) mutation in the Amyloid Precursor Protein (APP) is sufficient to cause early onset familial AD ^68^ and increased levels of Aβ_1-42_ ^69^, a variant of Aβ more likely to oligomerize ^70^ and to form disruptive plaques in the brain ^71^. Recently, increased levels of modified tubulin (polyglutamylated and/or Δ2) have been found in the hippocampi of postmortem patients with AD, suggesting that defects in α-tubulin re-tyrosination may be implicated in AD ^72^. Interestingly, fluctuations of detyrosinated tubulin in synaptosomal fractions from the dentate gyrus and corresponding MT instability/stability phases have been associated with associative learning and memory consolidation (Uchida et al., 2014). In that study, aged mice failed to regulate learning-dependent MT instability/stability phases and pharmacological disruption of either of the two phases led to deficits in memory formation. These data indicate that failure in regulating the Tyr/deTyr cycle occurs as a result of aging ^17^ and may play a primary role in synaptic plasticity and dementia related disorders. Moreover, oligomeric Aβ_1-42_ (oAβ) induces detyrosinated MTs in hippocampal neurons and this activity contributes to tau hyperphosphorylation and tau dependent synaptotoxicity ^73^. Finally, loss of MT dynamics was also reported in neurons from *Kif21b* KO mice that exhibit learning and memory disabilities ^74^. Despite these compelling evidences, whether perturbation of the Tyr/deTyr cycle is a molecular driver of synaptic pathology remains unexplored.

We hypothesized that loss of tubulin re-tyrosination and consequential accumulation of detyrosinated and Δ2 tubulin are molecular drivers of synaptic pathology by affecting MT dynamics in spines. Indeed, we found that in the hippocampus of TTL hemizygous (TTL^+/−^) mice, reduced levels of TTL expression led to significant changes in the Tyr/deTyr tubulin ratio and produced defects in synaptic transmission and plasticity that were associated with a loss of excitatory synapses. We examined whether TTL depletion was a *bona fide* feature of neurodegenerative disease and found that TTL was down-regulated in both sporadic and familial AD, and that abnormally high levels of detyrosinated and Δ2 tubulin accumulated in brain samples of AD patients. We explored whether TTL and dynamic MTs had a protective effect against the loss of synapses induced by oAβ. We found that MT entry into spines protected neurons from spine pruning and that acute oAβ exposure decreased the fraction of spines invaded by MTs prior to spine loss. Remarkably, TTL expression inhibited both spine loss and the decrease in the fraction of spines invaded by MTs, underscoring a role for re-tyrosinated tubulin in protecting synapses by driving dynamic MTs into spines.

Our data unveil a role for the Tyr/deTyr tubulin cycle in regulating cognitive parameters such as dendritic spine density, synaptic plasticity and memory. They also provide compelling evidence for dysfunction of the cycle in AD and suggest that regulation of α-tubulin re-tyrosination may be critical for shielding synapses against oAβ-induced synaptic injury by promoting invasion of dynamic MTs into spines.

## RESULTS

### Inhibition of tubulin re-tyrosination induces age-dependent synaptic defects

Dynamic MTs are crucial for synaptic plasticity and known to bear tyrosinated tubulin, and so we directly examined whether perturbation of the Tyr/deTyr tubulin cycle (Figure 1A) affects synaptic function. As total genetic ablation of TTL is perinatally lethal in mice ^61^, we used TTL hemizygous mice which are viable and fertile. Firstly, we confirmed that in protein extracts from hippocampi of 3- month-old TTL^+/−^ mice both TTL protein levels and Tyr/deTyr tubulin ratio were significantly reduced compared to wild-type (WT) mice (TTL = −44,95 ± 3,95 % of WT and Tyr/deTyr ratio = −39,46 ± 5.04 % of WT, Figure 1B-C and S1A-B).

**Figure 1.**
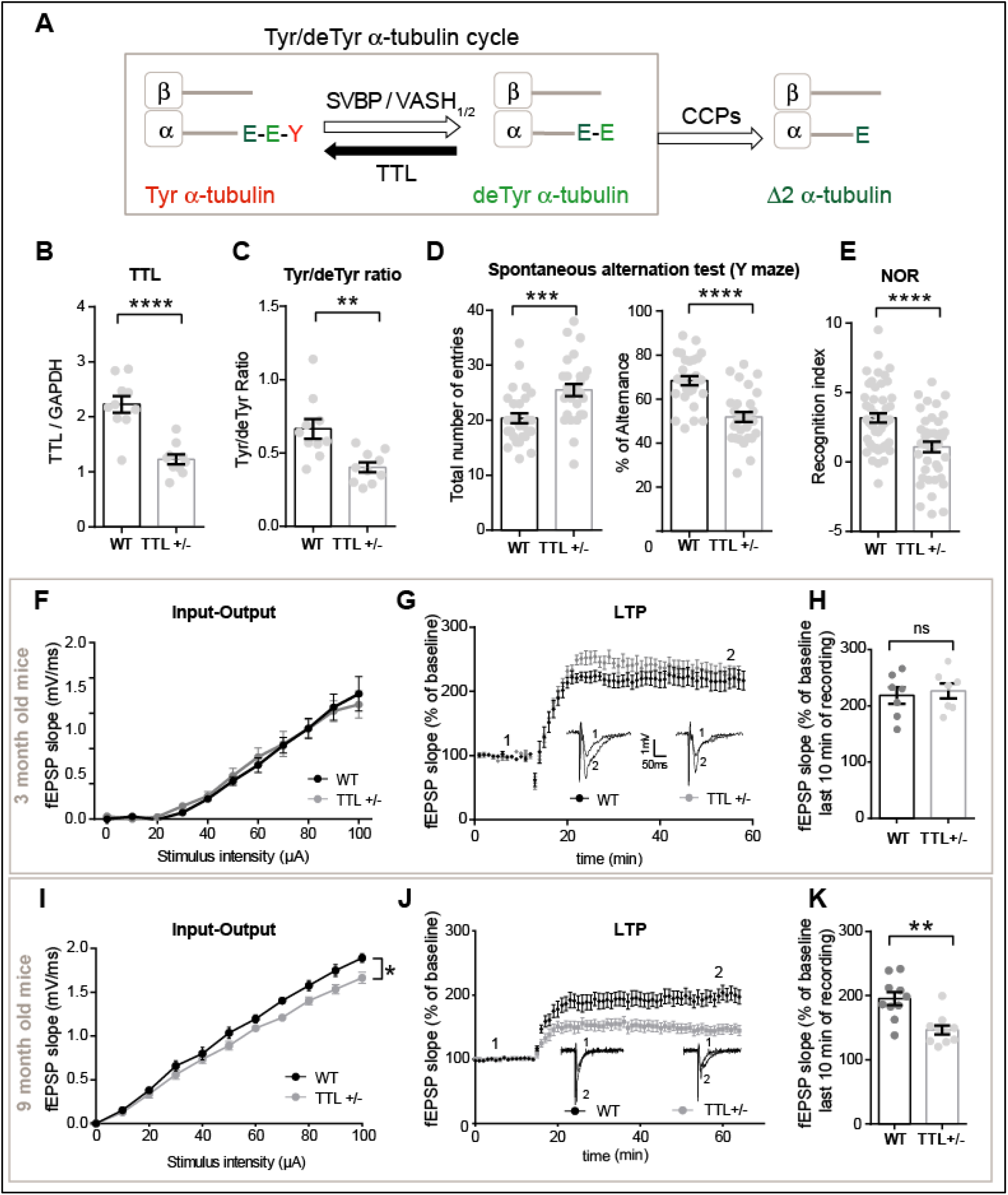
TTL reduction induces early memory defects and age-dependent alteration of synaptic plasticity. **(A)** Schematic representation of α-tubulin Tyr/deTyr cycle. TTL (tubulin tyrosine ligase), SVBP (small vasohibin-binding protein), VASH-1/2 (vasohibin-1 and −2), CCPs (cytosolic carboxypeptidases). **(B-C)** Relative amount of TTL (normalized with GAPDH) and Tyr/deTyr tubulin ratio, in protein extracts from hippocampi of 3-month-old WT and TTL^+/−^ mice. Graphs represents mean ± SEM. Mann-Whitney t test, ** p < 0.01, **** p < 0.0001. n = 10 independent experiments for each genotype. **(D)** Spontaneous alternation in Y-maze test. Total number of arm entries and percentage of alternance of 3-month-old WT and TTL^+/−^ mice. Graph represents mean ± SEM. n = 28 for WT and TTL^+/−^ mice. Student’s t test, *** p < 0.001, **** p < 0.0001. (**E**) Novel Object Recognition test. Recognition index (time spent exploring the novel object minus the time spent exploring the two familiar objects, in sec) of 3-month-old WT and TTL^+/−^ mice, measured 1h after familiarization. Mean ± SEM, n = 48 and 40 for WT and TTL^+/−^ mice, respectively. Student’s t test, **** p < 0.0001. (**F**) Input/output (I/O) curves of 3-month-old WT and TTL^+/−^ mice slices. Curves were constructed by plotting mean fEPSPs slopes ± SEM as a function of stimulation intensity. Two-way ANOVA, genotype x stimulation intensity interaction is not significant (F (10, 80) = 0,3845, p = 0.9500). n = 5 slices from 3 WT mice and n = 5 slices from 3 TTL^+/−^ mice. **(G)** Long-Term Potentiation (LTP) of 3-month-old WT and TTL^+/−^ mice. Curves represent normalized mean of fEPSPs slopes ± SEM as a function of time before and after LTP induction. Representative traces from one experiment are shown. They were extracted at the times indicated (1, 2) on the graph **(H)** Graph showing normalized mean of fEPSPs slopes ± SEM for the last 10 min of recording in WT and TTL^+/−^ mice. Mann-Whitney test, ns = not significant (p = 0.8048). n = 7 slices from 3 WT mice and n = 7 slices from 3 TTL^+/−^ mice. **(I)** Input/output (I/O) curves of 9-month-old WT and TTL +/− mice slices. Two Way ANOVA, genotype x stimulation intensity interaction (F (10, 220) = 1,923, * p = 0.0433). n = 12 slices from 5 WT mice and n = 12 slices from 5 TTL^+/−^ mice. **(J)** Long-Term Potentiation (LTP) of 9-month-old WT and TTL^+/−^ mice. Representative traces from one experiment are shown. They were extracted at the times indicated (1, 2) on the graph. **(K)** Graph showing normalized mean of fEPSPs slopes ± SEM for the last 10 min of recording in WT and TTL^+/−^ mice. Mann-Whitney test, ** p = 0.0021; n = 10 slices from 4 WT mice and n = 10 slices from 4 TTL^+/−^ mice.

TTL^+/−^ mice showed normal locomotor activities and sensorimotor functions as well as intact hippocampus-dependent spatial memory when assessed by the Morris Water Maze Test (Figure S2). These data are consistent with negligible effects of reduced Tyr/deTyr tubulin ratio on sensorimotor circuit development or hyperactivity and with the lack of manifested spatial navigation defects in most preclinical AD models at a young age ^75^. We also performed spontaneous alternation in Y-maze and novel object recognition tests (Figure 1D, E). These memory tests were selected because they broadly assess function of cognitive domains that correlate with neural circuitry disrupted early in AD, including the hippocampus ^76^, and have been useful to reveal memory defects in preclinical models of β-amyloidosis and tauopathy ^77, 78^. Indeed, TTL^+/−^ mice exhibited robust deficits in spontaneous alternation in Y-Maze (20.36 ± 0.91 versus 25.50 ± 1.09 number of entries and 68.44 ± 2.13 versus 51.88 ± 2.29 % of alternation for WT and TTL^+/−^ mice, respectively) (Figure 1D). Also, in the novel object recognition test, TTL^+/−^ mice spent significantly less time exploring the novel object than WT mice (delta between new and familiar object of 3.17 ± 0.33 versus 1.08 ± 0.37 sec for WT and TTL^+/−^ mice, respectively) (Figure 1E). Such behavioral deficits suggest impairments of spatial working and of short-term recognition memory.

Next, we investigated hippocampal synaptic transmission in 3 and 9-month-old WT and TTL^+/−^ mice. The efficacy of basal excitatory synaptic transmission was determined by field recordings of postsynaptic excitatory responses elicited by a range of electrical stimuli of axonal CA3-CA1 Schaffer collateral fibers, in hippocampal slices. While in 3-month-old animals, the input/output (I/O) curves revealed no differences between genotypes, in 9-month old mice, observed a significantly weaker postsynaptic response in TTL^+/−^ animals than in WT animals (Figure 1F, I) indicating defective basal synaptic transmission in older TTL^+/−^ mice. Furthermore, application of a theta-burst LTP protocol showed no difference in potentiation at 3-month-old mice between WT and TTL^+/−^ mice (Figure 1 G- H), but a reduced potentiation in 9-month-old TTL^+/−^ compared to WT mice (Figure 1 J-K, −25.04 ± 3.68 % of WT).

Altogether, these data demonstrate that a reduction in TTL expression results in loss of tyrosinated tubulin *in vivo*, early memory defects and age-dependent hippocampal synaptic dysfunction that affects both basal transmission and activity-dependent plasticity.

### Inhibition of tubulin re-tyrosination affects dendritic spine density

We examined the effects of TTL reduction at the level of individual neurons by measuring dendritic spine density and morphology both *in vivo* and using neurons in primary neuronal culture. Dendritic spines are often classified in three morphological types, corresponding to successive developmental stages: thin, stubby and mushroom-like spines ^79^. For *in vivo* evaluation, TTL^+/−^ deficient mice were crossed with Thy1-eYFP-H transgenic mice to visualize dendritic spines, and spine density evaluated in layer V cortical neurons ^80^. These neurons express moderate yellow fluorescent protein (YFP) levels, allowing accurate quantification of spine density (Figure 2 A), in contrast to hippocampal neurons in which expression levels were too high for proper assessment. Confocal microscopy of *in situ* cortical neurons from TTL^+/−^-Thy1-eYFP-H mice showed a 15.97 ± 2.6 % decrease in dendritic spine density compared to WT^+/−^-Thy1-eYFP-H littermates (2.147 ± 0.07 and 1.804 ± 0.05 spines/µm for WT and TTL^+/−^, respectively). The decrease mainly affected mature forms of dendritic spines (Figure 2 A-B). A comparable drop in mature spines (−15.53 ± 1.2 % of WT) was observed in cultured hippocampal neurons obtained from TTL^+/−^ embryos (1.204 ± 0.021 and 1.017 ± 0.014 spines/µm for WT and TTL^+/−^, respectively, Figure 2 C-D). Similar results were obtained when acute TTL knock-down was performed in rat hippocampal neurons using two independent TTL-targeting shRNAs (Figure 2E). TTL silencing resulted in an accumulation of Δ2 tubulin (Figure S1C) and induced a dramatic reduction of dendritic spine density (Figure 2E-F) with values similar to those observed in TTL KO neurons (Figure S3, −52.88 ± 2.67 %; −47.14 ± 4.30 % of WT for shRNA1, shRNA2 treated neurons respectively and −41.17 ± 1.25 % for TTL KO neurons). Together, these results show that reducing TTL expression affects the density of dendritic spines *in vitro* and *in vivo*, providing evidence for a novel role for tubulin re-tyrosination in regulating structural plasticity.

**Figure 2.**
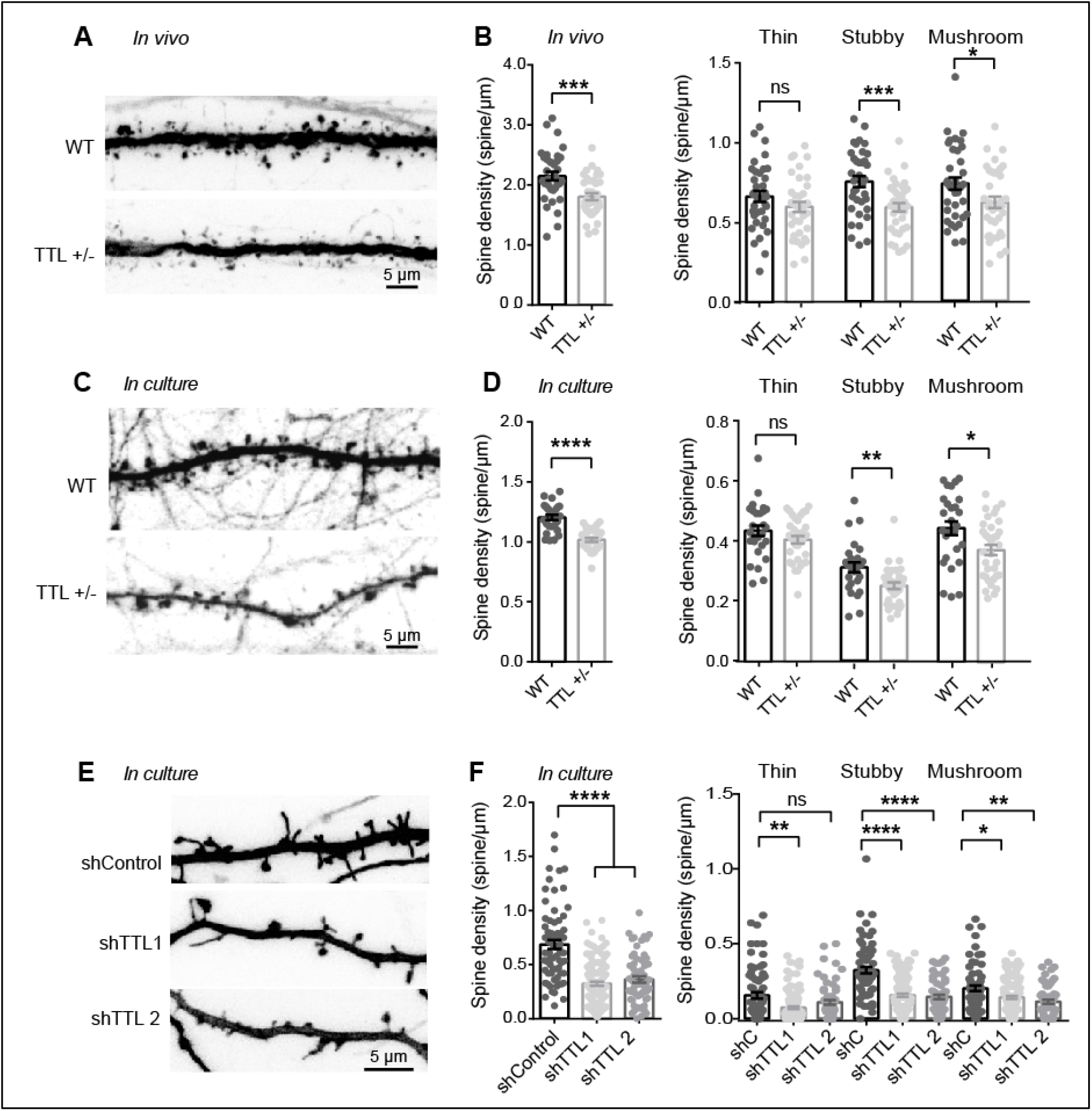
TTL reduction decreases dendritic spine density *in vivo* and in cultured neurons. **(A)** Confocal images showing representative examples of dendritic segments of cortical neurons from 4-month-old Thy1-eYFP-H WT and Thy1-eYFP-H TTL^+/−^ mice. **(B)** Total dendritic spine density, or that of each different morphological type of spines, is represented for Thy1-eYFP-H WT and Thy1- eYFP-H TTL^+/−^ cortical neurons. Graphs represent mean ± SEM. n = 36 neurons from 4 independent animals of each genotype. Student’s t test, * p < 0.05; *** p < 0.001 and ns = not significant. **(C)** Confocal images showing representative examples of the dendritic segments of GFP-expressing WT and TTL^+/−^ hippocampal neurons in culture at 17 DIV. **(D)** Total dendritic spine density, or that of each different morphological types of spines are represented for WT and TTL^+/−^ hippocampal cultured neurons. Graphs represent mean ± SEM. n = 27 and 34 neurons from WT and TTL^+/−^ embryos from at least 3 independent cultures. Student’s t test, * p < 0.05; ** p < 0.01; **** p < 0.0001 and ns = not significant. **(E)** Confocal images showing representative examples of dendritic segments of DiOilistic labeled WT rat hippocampal neurons in culture at 21 DIV, infected with control shRNA or shRNA targeting TTL (shTTL1 and shTTL2). **(F)** Total dendritic spine density or that of each different morphological types of spines, of hippocampal neurons infected with control shRNA (non-coding shRNA) or 2 independent shRNA lentiviruses targeting TTL (shTTL1 and shTTL2). Graphs represent mean ± SEM. n = 71, 124 and 60 neurons from control shRNA, shTTL1 and shTTL2 respectively, from at least 3 independent cultures. Kruskal-Wallis with Dunn’s multi-comparison test, * p < 0.05; ** p < 0.01; **** p < 0.0001 and ns = not significant.

### Tubulin re-tyrosination is perturbed in Alzheimer’s disease

The synaptic defects observed when levels of tyrosinated tubulin are perturbed raised the question whether dysregulation of tubulin re-tyrosination is a feature of AD, a neurodegenerative disorder in which synaptic pathology is prominent at early stages. We performed a detailed analysis of the relative amount of TTL, tyrosinated, detyrosinated and Δ2 tubulins in postmortem human brain tissues from sporadic AD patients and age-matched controls using ELISA and immunoblots. For these analyses, each AD brain was histologically analyzed according to Braak’s criteria ^81^ to discriminate early (Braak I-II), middle (Braak III-IV) and late AD stages (Braak V-VI), as shown in Table S1. AD sequentially affects the entorhinal cortex (E), hippocampus (H), temporal cortex (T) and lateral prefrontal cortex (L). We analyzed TTL levels and the different α-tubulin forms in protein extracts prepared from these four brain regions of AD patients and controls (Figure 3A-E). Global analysis indicated a statistically significant effect of Braak stages on TTL content (F (3, 25) = 4.3454, p = 0.0135, Figure 3B, red box). Post-hoc comparison of TTL content in control and AD brains showed a significant decrease in temporal and prefrontal cortex of AD patients (*p = 0.0322 and *p = 0.012, respectively for Braak V-VI versus controls, Figure 3B). No significant effect of brain region on TTL content was observed (F (3, 75) = 0.2185, p = 0.8833, Figure 3B, red box) suggesting that the TTL decrease observed in AD samples affects the whole brain. Regarding tyrosinated tubulin levels, a global analysis indicated that there was no significant dependence on Braak stage (p = 0.3556, Figure 3C, red box). The same analysis indicated an effect of the brain region (F (3, 75) = 3.1183, p = 0.03102, Figure 3C, red box) but pairwise post-hoc tests failed to support this conclusion (all p values > 0.05). For detyrosinated and Δ2 tubulin levels, the Braak stage had a global significant effect (F (3, 25) = 3.515, *p = 0.0297 and F (3, 25) = 5.877, **p = 0.0035 for detyrosinated and Δ2 Tub, respectively, Figure 3D-E, red boxes). Post-hoc comparison of Δ2 tubulin content in each brain region as a function of Braak stage, indicated an increase of the amount of Δ2 tubulin in all regions in AD samples, as compared to controls (Figure 3E, **p = 0.0018, p = 0.0584, *p = 0.0195 and *p = 0.0144 for entorhinal, hippocampus, temporal and lateral cortex, respectively, for Braak V-VI versus controls). Interestingly, the brain region had a global significant effect on both detyrosinated and Δ2 tubulin levels (Figure 3D-E, red boxes, F (3, 75) = 8.190, ****p = 0.00008 and F (3, 75) = 10.091, ****p = 0,00001 for detyrosinated and Δ2 Tub, respectively). This was confirmed by pairwise post-hoc comparisons which indicated for example that Δ2 tubulin content, in the enthorinal cortex, was significantly higher than in the hippocampus in AD brains (p = 0.3437, **p = 0.0082, p = 0.0572 and *p = 0.0127, for control, Braak I-II, Braak III-IV and Braak V-VI, respectively; pairwise t test with Sidak correction). Altogether, these results indicate that in AD conditions, a global TTL impairment is present from an early stage of the disease and is associated with increased amounts of non-tyrosinated tubulin especially in the entorhinal cortex, the first region to be affected in AD ^82^.

**Figure 3.**
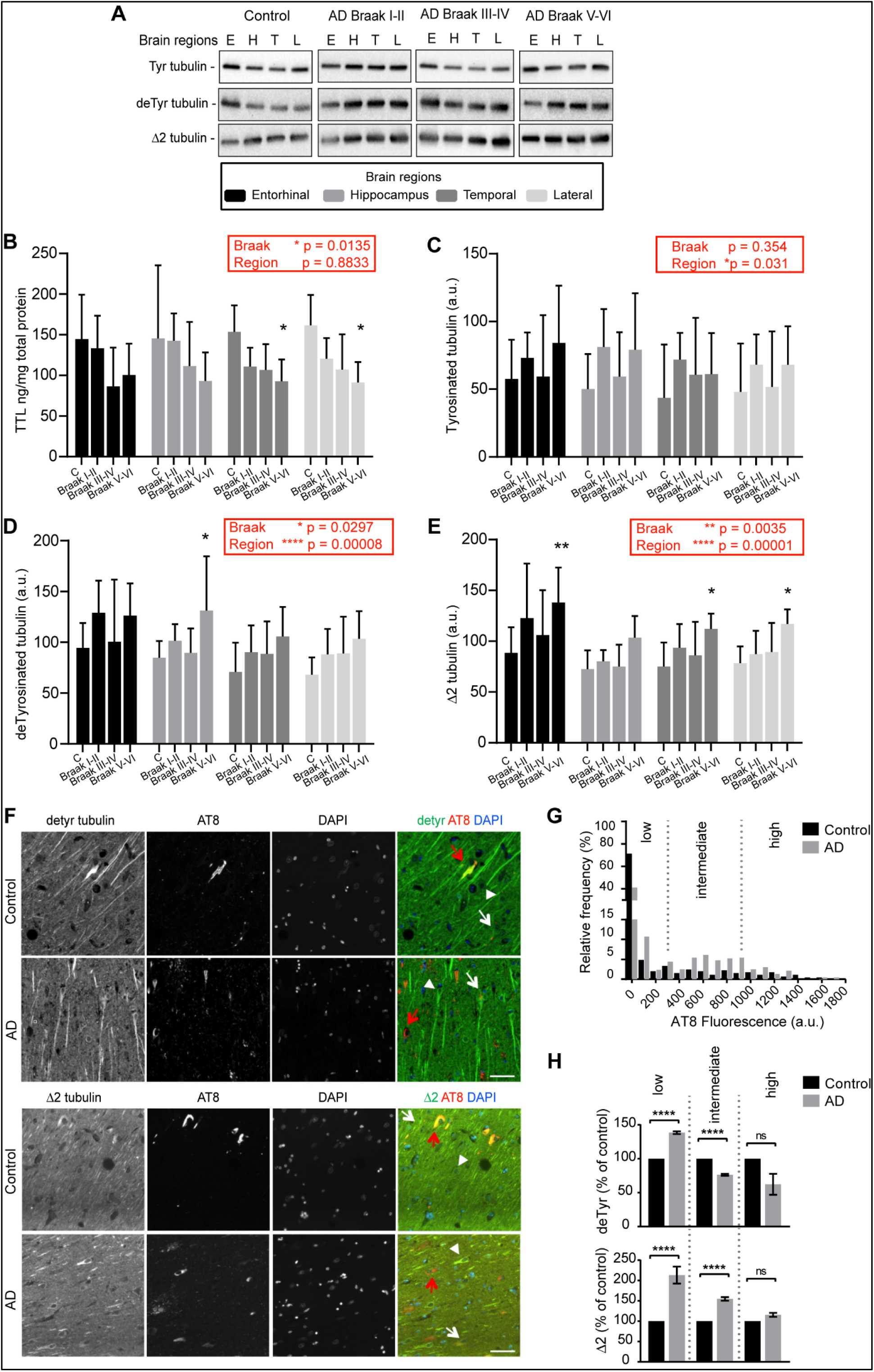
Loss of TTL and increased non-tyrosinated tubulin levels in sporadic AD brain samples. **(A)** Representative immunoblot analysis of tyrosinated, detyrosinated and Δ2 tubulin levels in brain homogenates from entorhinal cortex (E), hippocampus (H), temporal (T) and lateral prefrontal cortex (L) from control, early AD (Braak I-II), middle AD (Braak III-IV) and late AD (Braak V-VI) patients. **(B-E)** Quantification of TTL protein expression and modified tubulins levels (tyrosinated, detyrosinated and Δ2 tubulin) in each brain region from control and Alzheimer’s disease (AD) patients. Graphs represent mean ± SEM. The dependence of protein levels on, respectively, clinical stage and brain area was quantitated in each case using a linear mixed model, with Braak stage and brain region as fixed effect factors. Boxed p values measure the overall significance of these factors (type II Wald F test of model coefficients). In each brain area, post-hoc testing of variations due to individual Braak stages was performed by Dunnett’s test of differences with control. Significance levels are indicated as follows: * *p* < 0.05; ** *p* < 0.01; *** *p* < 0.001 and **** *p* < 0.0001. n = 11, 5, 6, and 7 for Control, Braak I-II, Braak III-IV and Braak V-VI AD patient brains, respectively. (**F**) Representative images of detyrosinated, Δ2 tubulin and phospho-tau in pyramidal neurons of hippocampi from AD patients. Dual immunostaining of detyrosinated (upper panel) or Δ2 tubulin (lower panel) and AT8-reactive phospho-tau, combined with nuclear staining with DAPI, was performed on sections of control and AD patient hippocampi. Neurons with low (white arrowheads), intermediate (white arrows) or high (red arrows) levels of AT8 immunofluorescence are shown. (**G**) Relative frequency distribution of phospho-tau (AT8) immunofluorescence levels (arbitrary units) in pyramidal neurons of control and AD brains. Low, intermediate, and high phospho-tau groups were defined based on fluorescence intensity. Two-sample Kolmogorov-Smirnov test, **** p < 0.0001. (**H**) Intensity of detyrosinated tubulin (upper graph) or Δ2 tubulin (lower graph) immunofluorescence in pyramidal cell bodies of AD hippocampal neurons relative to control, shown as a function of AT8 labelling level. Mean ± SEM. For deTyr tubulin, n = 382, 67 and 11 neurons in controls and n = 296, 162 and 2 for AD neurons in low, intermediate and high phospho-tau groups, respectively. For Δ2 tubulin, n = 249, 45 and 2 neurons in controls and n = 91, 133 and 102 for AD neurons in low, intermediate and high phospho-tau groups, respectively. Mann-Whitney test, ns = not significant and **** p < 0.0001.

We next analyzed modifications in non-tyrosinated tubulin content *in situ* by performing an immunocytochemistry study of AD brains. We performed a semi-quantitative immunofluorescence analysis of cell bodies and proximal dendrites of randomly selected individual pyramidal cells in the anterior hippocampal formation of sections from AD and controls (Table S2, Figure 3F-H). Each selected neuron was classified for tau pathology with either low, intermediate or high level of AT8 labelling (Ser202 and Thr205 phospho-tau antibody) and the mean intensity of detyrosinated and Δ2 tubulin staining was calculated. As expected, strongly AT8-reactive neurons were far more frequent in the AD samples, consistent with the pathological scoring of control and AD post-mortem human brains (Figure 3G, Table S2). Interestingly, we found that AD neurons with relatively low levels of phospho-tau, and thus presumably at an early stage of the degeneration process, were significantly enriched in detyrosinated and Δ2 tubulin compared to non-diseased neurons (Figure 3H, **** p < 0.0001). In contrast, AD neurons with the stronger AT8 staining displayed a lower level of detyrosinated tubulin than that of highly AT8-reactive neurons from control hippocampi, presumably as a consequence of advanced neurodegeneration and/or accelerated conversion of detyrosinated to Δ2 tubulin in diseased brains. These in situ results confirmed the accumulation of non-tyrosinated tubulin in pyramidal neurons in AD conditions and indicated it may occur at an early stage of the disease.

To explore whether perturbation of tubulin re-tyrosination and MT dynamics was a hallmark of familial AD, we utilized isogenic hiPSC lines in which the AD-linked London mutation (V717I) was knocked-in via CRISPR/Cas9 into one allele of the APP gene to replicate the genuine familial AD genotype ^83^. hiPSCs harboring the London mutation and the isogenic control parent line were differentiated *in vitro* into human cortical neurons via an neural progenitor (NPC) intermediate as previously described ^83, 84^. After 30 to 40 days of differentiation, a time at which differentiated cortical neurons establish synapses, neurons were lysed, and TTL, detyrosinated and Δ2 tubulin levels analyzed by immunoblot. At this stage of differentiation, the mutant neurons accumulated tau protein, which was phosphorylated (tau46 and AT8, Figure 4A-B), confirming the occurrence of a previously described pathological feature associated with this APP mutation ^69^. Consistent with our observations of brain samples, neurons with mutant APP displayed a significant reduction in TTL content (Figure 4C), an increase in Δ2 tubulin levels and showed a trend in the accumulation of detyrosinated tubulin compared to isogenic controls (Figure 4D-E).

**Figure 4.**
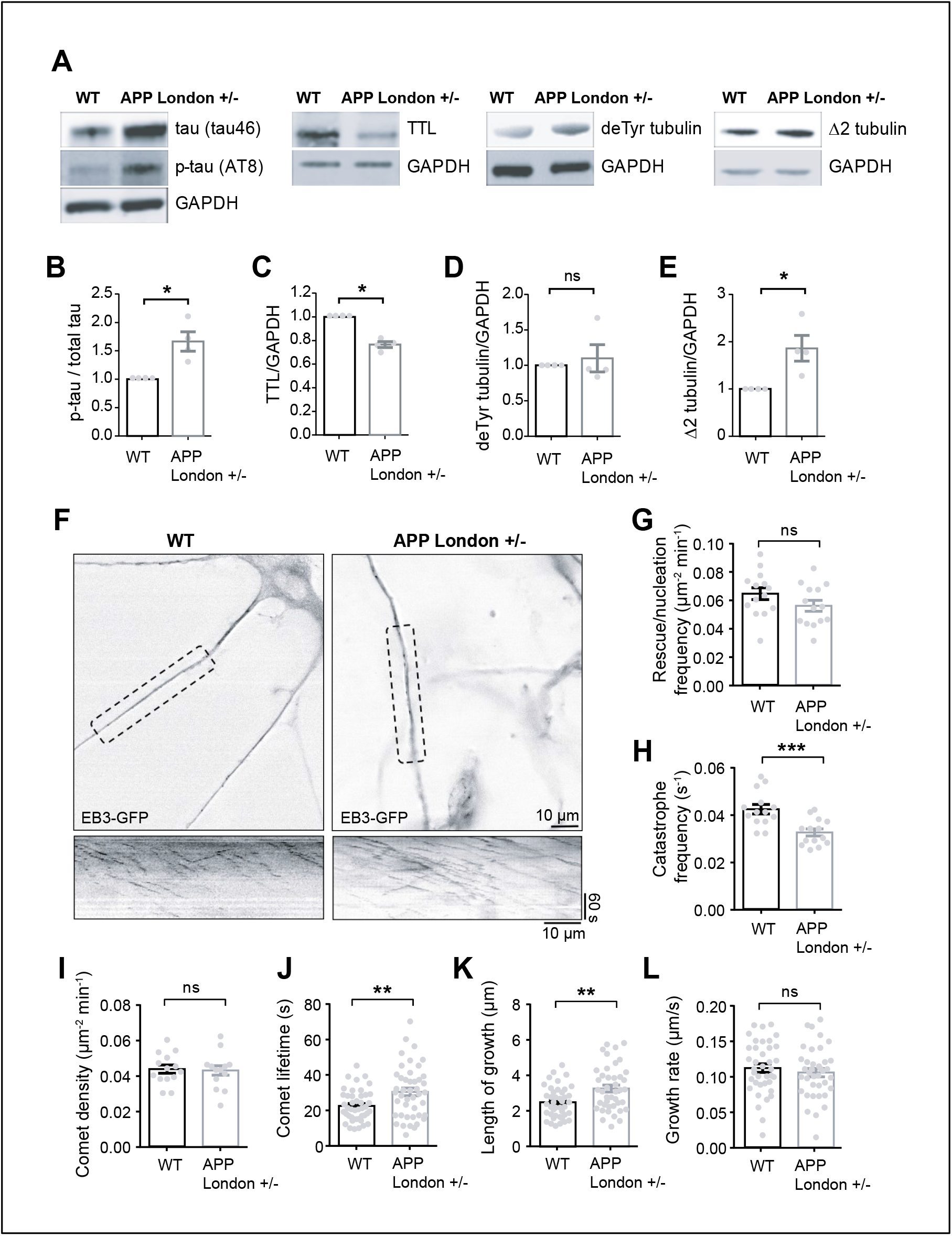
Loss of TTL and increased non-tyrosinated tubulin levels correlate with inhibition of MT dynamics in human cortical APP London neurons. **(A)** Human cortical neurons, derived from WT and APP London (V717I) knocked-in hiPSCs isogenic lines, were lysed between 30 and 40 days of differentiation and probed for phospho-specific tau (AT8), total tau (tau46), TTL, detyrosinated tubulin and Δ2 tubulin. GAPDH was used for normalization. Immunoblot quantifications of phospho-tau normalized to total tau **(B)**, TTL **(C)**, detyrosinated **(D)** and Δ2 tubulin **(E).** Data are expressed as a ratio of WT and graphs represent mean ± SEM. n = 4 independent neuronal differentiation experiments. Mann Whitney test, ns = not significant, * p < 0.05. **(F)** WT and APP London human cortical neurons expressing EB3-GFP. Representative neurites (dashed boxes) from human cortical neurons were analyzed for MT dynamics and kymographs of these regions are shown below. Scale bar: 10 μm. (**G-L**) Parameters of MT dynamics are represented as mean ± SEM. n = 14 neurites from WT and APP London neurons for **G** to **I**, and n = 44 comets for **J**, 42 comets for **K** and 38 comets for **L**, from WT and APP London neurons respectively. Student’s t-test, ns = not significant, ** p < 0.01 and *** p < 0.001.

We next directly examined MT dynamics in human neurons by transiently expressing EB3- eGFP to track the dynamic behavior of MT plus ends, and found that in neurons with APP mutation, while comet density and growth rates were not affected, catastrophe frequency, comet lifetime and length of growth were significantly reduced as compared to WT controls, a result consistent with inhibition of MT dynamics by resistance to undergo MT depolymerization (Figure 4G-M). Together, our results indicate that tubulin re-tyrosination is affected in sporadic and familial AD and that inhibition of MT dynamics observed in APP London human neurons is consistent with a disrupted Tyr/deTyr tubulin cycle in AD.

### Tubulin re-tyrosination protects neurons from oAβ synaptotoxicity and promotes MT invasion into spines

APP variants such as the London mutant generate larger amounts of amyloid beta peptide (Aβ) ^85^ and soluble oligomeric Aβ_1-42_ (oAβ) has been proposed to contribute to loss of synapses at an early stage of AD ^46^. We analyzed the consequences of the presence of oAβ, on cultured hippocampal neurons. First, we observed that neurons exposed to oAβ lost their spines in a time-dependent manner (−6.80 ± 4.63 %, −19.56 ± 4.41 %; −36.4 2 ± 2.79 % and −40.33 ± 6.57 % of control cells after 1h, 2 h, 3 h and 6 h of oAβ exposure, respectively) (Figure 5A-B). Next, we analyzed the dynamics of MTs invading into individual spines of neurons co-transfected with plasmids expressing EB3-eGFP and DsRed as a cell filler, in response to oAβ (Figure 5C). The dynamic parameters of spine-invading MTs (length of growth, comet lifetime, MT growth rate) and spine invasion lifetime were not affected by oAβ (Figure S4). However, oAβ acutely inhibited MT entry into spines at 0.5 h, while inducing a time-dependent renormalization of the fraction of MT-invaded spines starting at 2 h (3.68 ± 0.21 %, 1.03 ± 0.29%, 5.58 ± 0.54%, 4.97 ± 0.48 % and 4.70 ± 0.77% of spines for 0 h, 0.5 h, 2 h, 3h and 6 h of treatment respectively, Figure 5D), an effect possibly due to the reduction of the total number of spines over time (Figure 5B). We tracked and quantified the morphology of the same spines invaded or not invaded by MTs in neurons treated with vehicle or oAβ for 2 h (Figure 5E). In the absence of oAβ, MT-invaded thin spines appeared to switch more frequently to the larger stubby and mushroom spine types (Figure S5A), a phenotype in agreement with previous observations reporting modifications of spine morphology upon MT entry ^11^. However, in the presence of oAβ, spines that were not invaded by dynamic MTs had a higher chance of being pruned (Figure 5E & S5B) and the non-invaded mushroom spines that did not collapse showed increased transitions to stubby or thin spines, presumably causing additional loss of synaptic strength. For example, after 2h of oAβ treatment, only 9 % of MT-invaded spines were pruned compared to 35 % of non-targeted spines (Figure 5E).

**Figure 5.**
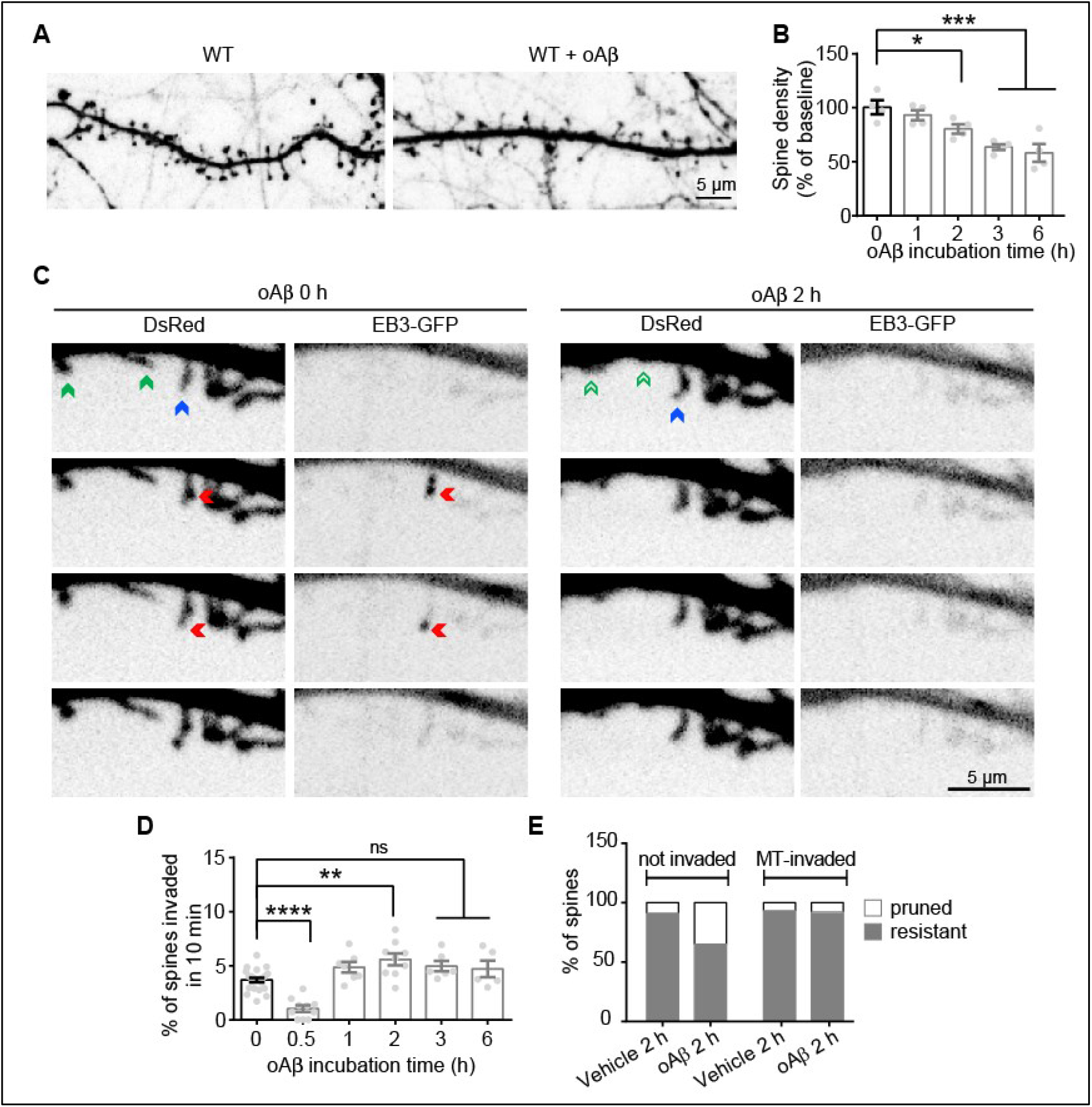
Acute oAβ treatment affects spine invasion by dynamic microtubules in cultured neurons. (**A**) Confocal images showing representative examples of dendritic segments of eGFP expressing WT rat hippocampal neurons (17 DIV) treated with DMSO or with 250 nM of Aβ oligomers for 2 days. (**B**) Graphs of the percentage of dendritic spine density in WT cultured neurons incubated with oAβ over 6 h. Data are expressed as a % of baseline and graphs represent mean ± SEM. n = 4 neurons analyzed over time. Repeated measures ANOVA with Holm-Sidak’s multiple comparison test, * p < 0.05 and *** p < 0.001. (**C**) Representative stills from videos of a WT neuron (21 DIV) transfected with DsRed and EB3-eGFP to visualize dendritic spines and the growing plus ends of MTs, before and 2 h after oAβ treatment. Spines that will prune are highlighted with a green arrow at time 0, and with an empty green arrow after 2h of oAβ treatment. The spine that will be invaded by MTs is highlighted with a blue arrow at time 0 and persists after 2h of oAβ treatment. MT invasion into the spine is highlighted with a red arrow. (**D**) Percentage of spines invaded by MTs before and after oAβ exposure at the indicated times. Graphs represent mean ± SEM. n = 22, 10, 9, 6 and 5 neurons at each time point. One-way ANOVA with Tukey’s multiple comparison test, ns = not significant, ** *p* < 0.01 and **** p < 0.0001. (**E**) Total percentage of spine pruning or resistance to vehicle or oAβ incubation. Graph represents the mean percentage of non-invaded spines (left) or MT-invaded spine fate (right). MT-invaded spines were significantly more resistant to oAβ-induced pruning than non-invaded spines (X^2^ = 43.64, 3 df, **** p < 0.0001, chi-square test).

These results indicate that oAβ causes early inhibition of MT entry into spines, and that these changes may be functionally related to the onset of spine pruning. The renormalization of the percentage of MT-invaded spines that we observed at later time points might reflect a relative accumulation of a class of spines which are intrinsically resistant to pruning. These results further suggest that entry of dynamic MTs, which are mainly composed of tyrosinated tubulin, may underlie the resistance of dendritic spines to synaptic injury by oAβ.

We examined the effect of chronic exposure to oAβ on TTL and tubulin tyrosination levels in primary cultured neurons. We found that a 48 h of oAβ exposure resulted in a 25.77 ± 5.23 % reduction in TTL content (Figure 6A), similar to what we observed in sporadic and familial AD samples (Figure 3B and Figure 4A, C). Lentivirus-driven TTL expression in these samples was performed to an extent that did not significantly affect Tyr/deTyr-tubulin ratio nor spine density in control neurons (Figure 6A-C) and we then tested for oAβ-induced spine pruning. Strikingly, in TTL-expressing neurons, oAβ completely failed to diminish spine density (Figure 6C-D), indicating that spine loss induced by oAβ might rely on downregulation of TTL and tyrosinated tubulin levels. While global biochemical analysis showed that oAβ did not appreciably alter the proportion of tyrosinated tubulin in these neurons (Figure 6B), it was conceivable that oAβ might have locally affected the pool of tyrosinated, dynamic MTs available for spine entry. To explore this possibility, we set out experiments to examine whether the percentage of spines invaded by dynamic MTs correlated with spine resistance to oAβ in neurons ectopically expressing TTL (Figure 6E-G). We found that expression of TTL averted the oAβ-induced drop in spine invasions by dynamic MTs measured at 30 min (Figure 6F) and oAβ-promoted spine loss which became detectable only 2.5 h later (Figure 6G).

**Figure 6.**
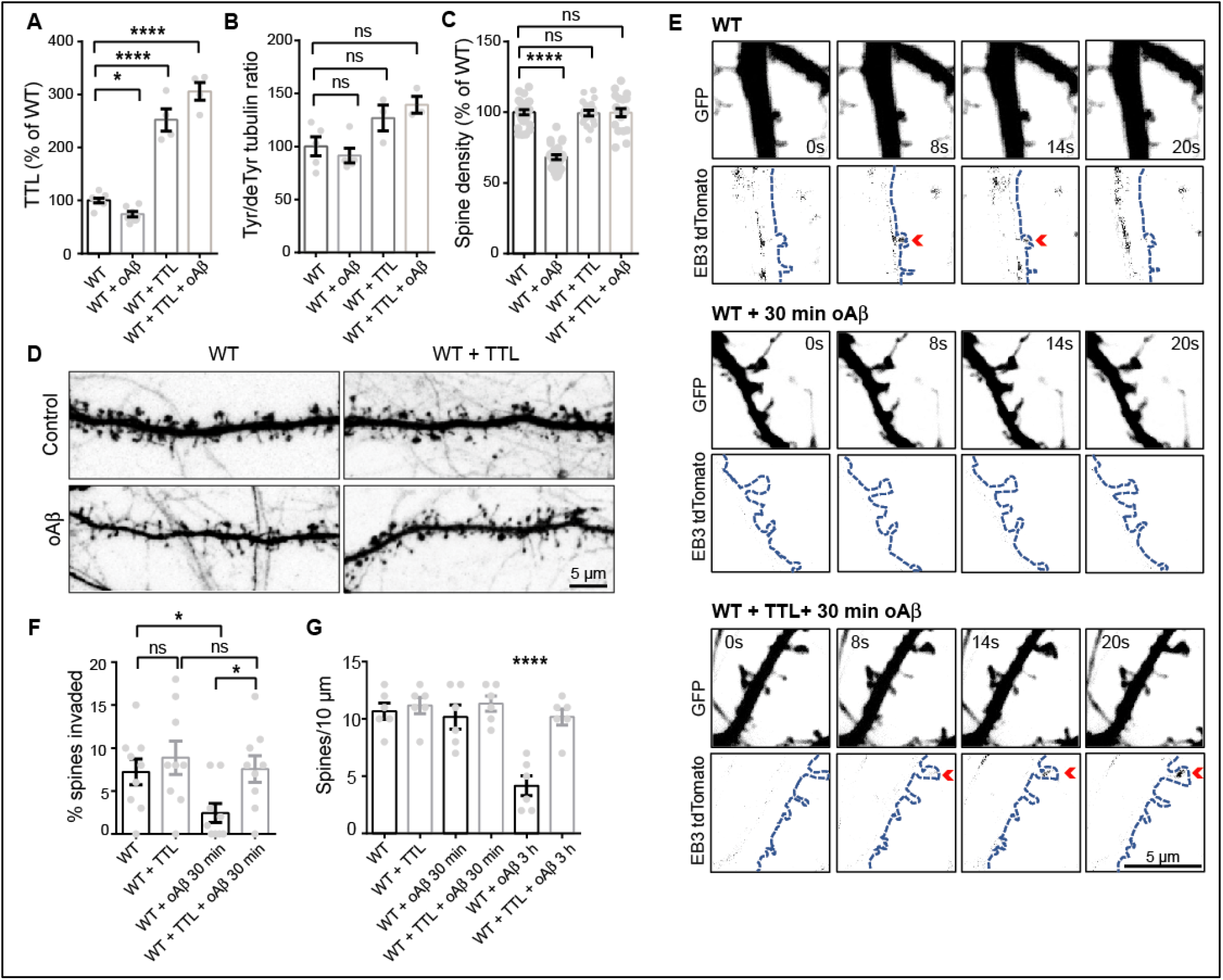
Ectopic TTL expression rescues neurons from oAβ-induced dendritic spine loss and resumes MT invasions into spines. Immunoblot analysis of TTL (**A**) and Tyr/deTyr tubulin ratio (**B**) from WT mouse cortical neurons (17 DIV) transduced or not with a lentivirus expressing TTL and chronically treated with DMSO or with 100 nM oAβ. Data are expressed as a % of WT and graphs represent mean ± SEM. (**A**) n = 8, 7, 4 and 4 cultures for WT, WT+ Aβ, WT+TTL and WT+Aβ+TTL respectively. Two Way ANOVA, oAβ treatment x TTL expression interaction (F (1, 19) = 14.6, ** p = 0.0012). All values were compared to WT, Holm-Sidak’s multiple comparison test, * p<0.05 and **** p<0.0001. (**B**) n = 5, 5, 3 and 3 cultures for WT, WT+ Aβ, WT+TTL and WT+Aβ+TTL respectively. Two Way ANOVA, oAβ treatment x TTL expression interaction (F (1, 12) = 1.309, p = 0.274). All values were compared to WT, Holm-Sidak’s multiple comparison test, ns = not significant. (**C**) Graphs of total dendritic spine density in cultured WT neurons treated as in A. Graphs represent mean ± SEM. n = 27, 26, 20 and 20 neurons for WT, WT+ Aβ, WT+TTL and WT+Aβ+TTL respectively. Two Way ANOVA, oAβ treatment x TTL expression interaction (F (1, 89) = 58.44, **** p < 0.0001). All values were compared to WT, Holm-Sidak’s multiple comparison test, ns = not significant and **** p<0.0001. (**D**) Confocal images showing representative examples of dendritic segments of GFP-expressing WT hippocampal mouse neurons (17 DIV) chronically treated with DMSO or with 100 nM oAβ. (**E**) Representative stills from videos of rat WT neurons (18 to 21 DIV) transduced or not with a TTL containing lentivirus and transfected with plasmids encoding eGFP and EB3-tdTomato to visualize the dendrites and spines and the growing plus ends of MTs, respectively. Cells were incubated with vehicle or with 250 nM of oAβ for 30 min. MT invasions of spines are highlighted with a red arrow. (**F**) Percentage of spines invaded by MTs after vehicle or oAβ exposure. Graphs represent mean ± SEM. n = 9 neurons for each condition. Two Way ANOVA, oAβ treatment x TTL expression interaction (F (1, 32) = 1.299, p = 0.275). Uncorrected Fisher’s Least Significant Difference, ns = not significant, * p < 0.05. (**G**) Graphs of total dendritic spine density in cultured neurons treated as in E and incubated with vehicle or with oAβ for 30 min or 3 h. Graphs represent mean ± SEM. n = 6 neurons of each condition. One-way ANOVA with uncorrected Fisher’s Least Significant Difference, ns = not significant, **** p < 0.0001.

Together, our results indicate that entry of dynamic tyrosinated MTs into spines underlie enhanced resistance of dendritic spines to synaptic injury and that restoring TTL expression can protect dendritic spines from oAβ toxicity.

## Discussion

In this study, we identify a role for the re-tyrosination of α-tubulin by TTL activity in the maintenance of synaptic function and AD-related synaptic dysfunction.

Our biochemical and immuno-histological analysis of TTL^+/−^ mouse hippocampi confirmed that hemizygous suppression of TTL leads to 40% reduction of tyrosinated tubulin at 3 months, and that this reduction is compatible with viability and normal life span. This result suggests that TTL levels are rate-limiting for the maintenance of physiological amounts of tyrosinated tubulin *in vivo*. We found that the behavioral performance of TTL^+/−^ mice at 3 months revealed impairments in spontaneous alternation test and novel object recognition but no defect in spatial learning assessed by Morris Water Maze, the standard test for evaluating hippocampal-dependent memory in rodents. This behavioral profile was consistent with no alteration in hippocampal basal transmission and CA3/CA1 LTP at this early age, which was instead characterized by deficits in spatial working and intermediate-term recognition memory most likely caused by cortical circuitry dysfunction. In agreement with synaptic cortical damage at this age, we observed loss of dendritic spines in serial sections obtained from cortical layer V of 3-month-old TTL^+/−^ mice. At 9 months, however, TTL^+/−^ mice had a clear reduction in their basal hippocampal transmission, a defect consistent with decreased spine density observed in cultured hippocampal neurons from TTL^+/−^ mice or transiently silenced of TTL expression. In addition, a striking decline in the LTP of synaptic strength at the Schaffer collateral synapses was observable in 9-month-old TTL^+/−^ mice, demonstrating that TTL deficiency exacerbates synaptic plasticity defects with aging.

Our in vitro analyses strongly suggest that these alterations may be related to defects in synaptic MT dynamics. In support of this model, we found that entry of dynamic MTs into spines correlated with resistance to oAβ-induced spine pruning. In addition, expression of TTL, in oAβ- treated neurons, prevented both transient loss of MT entry into spines and spine pruning, indicating that restoring MT invasions into spines is the mechanism by which TTL antagonizes oAβ-induced loss of synapses. We noticed that in the absence of oAβ, MT invaded spines tend to display higher structural plasticity, particularly in thin spines switching to larger headed spine types, which is in keeping with changes in spine head size associated with MT spine invasion. ^10, 11^. The fine-tuning of the Tyr/deTyr tubulin cycle as a function of local cues may be especially important in the vicinity of synapses which are particularly dependent on entry of dynamic tyrosinated MTs ^10, 11^. Live imaging of spines invaded by MTs during incubation with oAβ showed that the minority of spines that were invaded by MTs during the recording period had a fate that clearly differed from that of neighboring, non-invaded spines, strongly supporting the notion that MT entry is critical for the delivery or removal of cargos that regulate spine stability and plasticity. Given the pleiotropic effects that the Tyr/deTyr cycle plays in the regulation of specific tracks for neuronal transport ^45, 86–88^, re-tyrosination of tubulin by TTL might be critical for the local recruitment or removal of spine modulating cargos specifically trafficked along tyrosinated MTs. Altogether, the electrophysiological, spine density and behavioral profile of TTL hemizygous mice shows that TTL is required for synaptic maintenance and plasticity, and that TTL deficiency increases synaptic vulnerability. These findings are relevant to the onset of synaptic dysfunction in neurodegenerative disease, as we find that TTL is down-regulated in AD brain, human AD neurons, and primary neurons exposed to oAβ. Biochemical analysis of post mortem brain samples from clinically graded AD patients indicated a robust loss of TTL and a gain in detyrosinated and Δ2 tubulin compared to samples from non-affected individuals in the same age range. The correlation between disease conditions and non-tyrosinated tubulin accumulation was confirmed at the single neuron level by imaging analysis of AD hippocampal sections. Deficits were narrowed to an early phase of the disease, a stage at which neuron morphology appears normal with deficiencies mainly affecting the synaptic compartments. The finding that the knock-in of the AD-linked London mutation in APP in *in vitro* differentiated human neurons also resulted in a drop in TTL compared to isogenic controls strongly supports a causal relationship between TTL loss and familial AD. Because the London mutation leads to an increase in the amyloidogenic processing of APP and overproduction of toxic Aβ species ^85^, the finding suggests that TTL down-regulation could be initiated by either defective APP processing and/or accumulation of oAβ. Indeed, chronic incubation of cultured mouse neurons with synthetic oAβ elicited a significant decline in TTL levels, although the underlying mechanisms are yet to be defined. The altered synaptic phenotype of TTL^+/−^ mice suggests that down-regulation of TTL might in turn aggravate oAβ synaptoxicity by reducing MT dynamics, and thus cause further loss of synapses. This notion would be consistent with the protection against dendritic spine retraction that we observed in neurons in which TTL was ectopically expressed.

Altogether, our results point to a modulatory role of the Tyr/deTyr tubulin cycle in synaptic plasticity and indicate that loss of TTL and tubulin re-tyrosination are features of AD that could play pathogenic roles at an early stage of the disease. The results also clearly indicate that in the early stages of AD, the MT network appears to be less dynamic than in normal conditions, with critical loss of dynamic MTs. They also suggest that the decrease in dynamic MTs, rather than a global MT destabilization, initiates AD pathology. Our pathogenesis model does not reject loss of MT integrity as a major pathological feature of advanced AD, but rather proposes that amyloidogenic APP processing may affect synaptic function by reducing the population of dynamic MTs entering into synapses at an early stage of the disease. While the molecular factors associated with the resistance of dynamic MT invaded spines remain to be identified, our results suggest that TTL activators may be beneficial to restore circuit integrity in sporadic and familial AD. In addition, vasohibins/SVBP carboxypeptidase (TCP) has been recently identified as a tubulin detyrosinating complex ^20, 21^, suggesting that also drugs able to modulate TCP activity may offer a valuable new approach for therapeutic intervention in AD.

## Supporting information

Supplemtary Information Peris et al

## Acknowledgements

We thank F. Vossier and L. Macedo for technical support; S. Andrieu, L. Romian, F. Mehr, F. Rimet, and S Bama-Toupet for animal care; Ju. Brocard for help in lentivirus preparation and E. Tein for conducting hiPSC neuronal differentiations. We are grateful to J. Crary (Icahn School of Medicine at Mount Sinai) and the Pathology Dept at CUIMC for kindly providing AD and age matched control brain slices from *post-mortem* tissue. We thank M. Gagliardini and S. Jules for helping with the analysis of *post-mortem* specimens and S. Pierre-Ferrer for the characterization of lentiviral shTTL vectors in primary neuronal cultures. Part of the work was performed at Grenoble Institut Neuroscience Photonic Imaging Center (part of the IBiSA-accredited ISdV core facility) and in CEA- IRIG animal facility (GRAL, ANR-17-EURE-0003). This work was supported by INSERM; CEA; CNRS; University Grenoble Alpes; France Alzheimer (CAPAlz-AAP SM 2018) and ANR (SPEED-Y, ANR-20-CE16-0021) grants to MJM; NIH/NIA RO3 AG060025, NIH/NIA RO1 AG050658 and NIH/NINDS R21 NS120076-01 grants to FB; RO3 AG060025 (Co-I), the Henry and Marilyn Taub Foundation, and the Thompson Foundation (TAME-AD) grants to AS. JP fellowship was from the Italian Academy at Columbia University and the Alzheimer’s Association Grant AARF-20-685875; MBR postdoctoral fellowship from Ramón Areces Foundation and salaries of JMH, CC and GF from a collaborative program between Servier laboratories and AA’s team.

## Contributions

LP, XQ, MJM, FB and AA conceived and designed the study. CB, GF, MEP, XQ, AK and JP performed molecular biology experiments and TTL lentivirus preparations. AA and MJM supervised TTL mice production, BDC and PD supervised behavioral tests and mouse brains biochemical studies. FL, FP and AB performed electrophysiological experiments. SGF performed and analyzed in vivo dendritic spine density. LP and JMH performed experiments to analyze spine density in mice hippocampal neurons. FB, MEP and JP designed and performed the analysis of spine density in rat hippocampal neurons. JMS and CC performed biochemical experiments with AD patient samples, JB and YG performed the associated statistical analysis. FB and XQ designed and performed in situ analysis of patient data. FB, XQ, JP and AK designed and performed analysis of microtubule entry into dendritic spines. FB, AK, MBR, JP and AS designed and performed experiments in human cortical neurons. LP, JP, YG, MJM, FB and AA wrote the manuscript, with contributions from all co- authors.

## MATERIAL AND METHODS

### Animals

All experiments involving mice were conducted in accordance with the policy of the Institut des Neurosciences de Grenoble (GIN) and in compliance with the French legislation and European Union Directive of 22 September 2010 (2010/63/UE). TTL heterozygous (TTL^+/−^) mice were obtained as previously described ^61^ and maintained in a C57BL6 genetic background by recurrent back- crosses with C57BL6 animals from Charles River Laboratories. Th1-eYFP line H mice ^80^ were obtained from Jackson Labs (B6.Cg-Tgn (Thy-YFP-H) 2Jrs) and crossed with TTL^+/−^ mice to generate a colony of C57BL6/Thy1-eYFP TTL mice. All experiments involving rats were approved by the Committee on the Ethics of Animal Experiments of Columbia University and performed according to Guide for the Care and Use of Laboratory Animals distributed by the National Institutes of Health. E18 pregnant Sprague Dawley rats were purchased from Charles River Laboratories.

### Electrophysiology

Electrophysiological tests were done with 3 and 9-month old WT and TTL^+/−^ mice. <u>Ex vivo slice preparation:</u> brain slices were prepared from 3 and 9-month-old WT and TTL^+/−^ mice. The brain was removed quickly and 350 μm thick sagittal slices containing both cortex and hippocampus were cut in ice-cold sucrose solution (in mM: KCl 2.5, NaH2PO4 1.25, MgSO4 10, CaCl2 0.5, NaHCO3 26, Sucrose 234, and Glucose 11, saturated with 95% O2 and 5% CO2) with a Leica VT1200 blade microtome (Leica Microsystemes, Nanterre, France). After the cutting, hippocampus was extracted from the slice and transferred in oxygenated ACSF (in mM: NaCl 119, KCl 2.5, NaH2PO4 1.25, MgSO4 1.3, CaCl2 2.5, NaHCO3 26, and Glucose 11) at 37 ± 1 °C for 30 minutes and then kept at room temperature for at least 1 hour before recordings. <u>Electrophysiological recordings</u>: each slice was individually transferred to a submersion-type recording chamber and continuously superfused (2ml/min) with oxygenated ACSF at 28°C. Extracellular recordings were obtained from the apical dendritic layers of the hippocampal CA1 area, using glass micropipettes filled with ACSF. Field excitatory postsynaptic potentials (fEPSPs) were evoked by the electrical stimulation of Schaeffer collaterals afferent to CA1. The magnitude of the fEPSPs was determined by measuring their slope. Signals were acquired using a double EPC 10 Amplifier (HEKA Elektronik Dr. Schulze GmbH, Germany), recorded with Patchmaster software (HEKA Elektronik Dr. Schulze GmbH, Germany) and analyzed with Fitmaster software (HEKA Elektronik Dr. Schulze GmbH, Germany). Input/output (I/O) curves characterizing basal glutamatergic transmission at CA3-CA1 synapses of WT and TTL^+/−^ mice were constructed by plotting mean fEPSPs slopes ± SEM as a function of stimulation intensity (10 to 100 μA). For Long-term potentiation (LTP) experiments test stimuli were delivered once every 15s and the stimulus intensity was adjusted to produce 40-50% of the maximal response. LTP was induced using a Theta Burst Stimulation (TBS involved 5 trains with 10 bursts of 4 pulses delivered at 100 Hz, an interburst interval of 200 ms and 20 s interval between each train). Average value of fEPSP slope was expressed as a percentage of the baseline response ± SEM.

### Behavioral studies

Behavioral tests were done in 3 to 4-month old WT and TTL^+/−^ mice. Evaluation of cognitive function was performed with spontaneous alternation in the Y maze test for working memory, and with the Novel Object Recognition test for episodic memory. <u>Spontaneous alternation test</u> was conducted in a Y-shaped maze, made of black Plexiglas. The maze is heightened approximately 1 meter high, constituted by 3 arms of equivalent size (L = 38 cm; W = 8 cm; H of walls = 15 cm), numbered from 1 to 3, and by equivalent angles between them (120°). The mouse was put in the center of the maze, the nose in the direction of the bottom of one of the arms. The mouse was free to explore the environment for 5 minutes. The experimenter observed the behavior by using a camera located in an independent room and noted the sequence of successive arm visits. A visit or an entrance into an arm was defined as four legs in the zone of the arm. The apparatus was cleaned with alcohol and subsequently with water between each mouse. An alternation is defined as a visit in a given arm followed by a visit into the other arm. The successive sequence of visits during 5 minutes determines the level of alternation. The performance of the animal is estimated by calculating a percentage of alternation: Alternation index = [number of alternations/ (total number of visited zones-2)] x 100. <u>Novel object recognition test</u> was performed in Y shaped maze, to about 1 meter in height, consisting of three opaque black plastic arms of equal size (L = 38 cm, W = 8 cm, H of wall = 15 cm), numbered 1 to 3, and at a 120° angle from each other. Four different objects by size, shape and pattern were used. The recognition test had three phases: habituation, familiarization and recognition. For habituation at day 1, the mouse was placed in the center of the Y-maze, without object, to freely explore the three arms for 10 min. For familiarization at day 2, the mouse was again placed in the center of the Y-maze which contained at each end different objects. The mouse freely explored for 5 min, during which it can familiarize with these three objects. For recognition test, 1h after familiarization, the mouse was placed in the center of the Y-maze where one object presented during the familiarization phase was replaced by a new object. The mouse freely explored for 5 min and the experimenter measured the time of exploration of each object using a semi-automatic key. The assessment criterion was the difference between the time of exploration of the new object and the mean time of the time of exploration of the two familiar objects. Recognition index = difference [New object – (Mean of the two familiar objects)] durations (in sec) of exploration.

### Plasmids

For lentiviral experiments, vector eGFP-pWPT (Addgene #12255, kind gift from D. Trono) was used to express eGFP, and cDNA encoding human TTL (NP_714923, Origene #RC207805L2) was cloned in it for TTL expression. PCR amplification and cloning of TTL cDNA was performed with Phusion DNA polymerase (Thermo Scientific) and In-Fusion HD Cloning kit (Clontech), respectively. eGFP cDNA was removed during the cloning process to produce an untagged TTL. For lentiviral shRNA expression, 2 TTL shRNA sequences, cloned in pLKO.1 vector, were purchased from Sigma-Aldrich: shTTL1 (TRCN0000191515, sequence: 5’ - CCG GCA TTC AGA AA GAG TAC TCA ACT CGA GTT GAC TAC TCT TTC TGA ATG CTT TTT TG - 3’) and shTTL2 (TRCN0000191227, sequence: 5’ - CCG GCT CAA AGA ACT ATG GGA AAT ACT CGA GTA TTT CCC ATA GTT CTT TGA GTT TTT TG - 3’) ^35^. The SHC001 pLKO.1-puro Empty Vector (Sigma) was used as control (shControl). For the transfection experiments, the plasmid encoding pCMV-EB3-EGFP was created by PCR of eb3 from mouse cDNA and insertion 3’ to pEGFP-C1 (kind gift from Dr. Frank Polleux). Kind gifts from Dr. E Dent include the plasmids EB3 tdTomato (Addgene #50708) and the plasmid encoding DsRed2 (Clontech), cloned into a pCAX vector. The plasmid pEGFP-N1 with a CMV promoter was also used (Addgene #6085-1). All constructs were verified by sequencing (Eurofins and Genewiz). Plasmids were purified with HiPure Plasmid Maxiprep kits (Invitrogen).

### Lentivirus production

Lentiviral particles were produced using the second-generation packaging system as previously described ^73, 89^. Lentivirus encoding GFP or TTL cDNA (packaging vectors, pWPT-based vector, Addgene, Cambridge, MA) and shTTL1, shTTL2 and control shRNA (packaging vectors pLP1, pLP2, and pLP-VSV-G, Thermofisher) were produced by co-transfection with the psPAX2 and pCMV-VSV-G helper plasmids, into HEK293T cells obtained from ATCC (ATCC-CRL-3216) using the calcium phosphate transfection method. Viral particles were collected 48 h after transfection by ultra-speed centrifugation, prior to aliquoting and storage at −80°C.

### Primary hippocampal neuronal cultures

Mouse hippocampi (E18.5) were digested in 0.25% trypsin in Hanks’ balanced salt solution (HBSS, Invitrogen, France) at 37°C for 15. min. After manual dissociation, cells were plated at a concentration of 5,000-15,000 cells/cm^2^ on 1 mg/ml poly-L-lysine-coated coverslips for fixed samples, or on ibidi glass bottom 60 µDishes for live imaging. Neurons were incubated 2 h in DMEM-10% horse serum and then changed to MACS neuro medium (Miltenyl Biotec) with B27 supplement (Invitrogen, France). Rat hippocampi were dissected from E18 embryos, and neurons were plated on 100 µg/ml poly-d-lysine–coated 12-well plates at the density of 3 × 10^5^ cells/well for biochemistry assays, 5 × 10^4^ cells/dish for live imaging in the chamber of 35- mm MatTek dishes or 4 × 10^4^ cells/coverslip on 18-mm coverslips for fixed samples. Primary neurons were maintained in Neurobasal medium (Invitrogen) with the supplement of 2% B-27 (Invitrogen) and 0.5 mM glutamine (Invitrogen), and one third of medium was changed every 3–4 d up to 4 weeks in culture.

### Lentivirus infection

To perform dendritic spine quantification, 1/100 of a hippocampal cell suspension was infected by 15 min incubation with GFP lentivirus (Lv) at a multiplicity of infection of 40. The infected population was then mixed with non-transduced cells before plating. Some of those cultures were infected at 1 day *in vitro* (DIV) with TTL lentivirus at a multiplicity of infection of 5. Hippocampal neurons were incubated for 18 DIV at 37°C, 5% CO2 in a humidified incubator and then fixed with 4% paraformaldehyde in 4% sucrose-containing PBS for 20 min. To induce acute TTL reduction, hippocampal neurons from WT rat embryos were infected at DIV 14 with lentiviral vectors containing either control or one of two independent TTL-targeting shRNAs and incubated until DIV 21. Ectopic expression of TTL for MT spine dynamics experiments was also achieved through lentiviral infection, with infection again occurring at DIV 14 and incubation until DIV21.

### Imaging of dendritic spines

For *in vivo* fixed samples, serial sections were obtained from cortical layer V of 3-month-old Thy1eYFP-H WT and Thy1eYFP-H TTL^+/−^ mice. For cultured samples, hippocampal neurons from WT, TTL ^+/−^ and TTL ^-/-^ embryos were infected with eGFP containing lentivirus and fixed at DIV18. Dendritic segments visualized by soluble eYFP and eGFP respectively, were obtained using a confocal laser scanning microscope (Zeiss, LSM 710). Serial optical sections (1024 × 1024 pixels) with pixel dimensions of 0.083 × 0.083 μm were collected at 200 nm intervals, using a × 63 oil-immersion objective (NA 1.4). The confocal stacks were then deconvolved with AutoDeblur. For in vitro analysis of spines in cultured hippocampal neurons isolated from rat embryos and infected with TTL-targeting shRNAs, DiOlistic labeling using the Helios gene gun system (Bio-Rad) was performed according to the manufacturer’s instructions. Tungsten particles (1.1 μm, Bio-Rad) coated with DiI (Invitrogen), which defines the neuronal architecture in red, were delivered into hippocampal neurons fixed in 4% PFA prior to mounting with ProLong Gold antifade mounting reagent (Invitrogen). Neurons were imaged the next day using an Olympus IX8Andor Revolution XD Spinning Disk Confocal System. Z stack images were taken at 0.2 um step length for 10-15 stacks and shown as maximum projections. Dendritic spine analysis (spine counting and shape classification) was performed on the deconvolved stacks using Neuronstudio and Neurolucida 360 ^90^. All spine measurements were performed in 3D from the z-stacks. The linear density was calculated by dividing the total number of spines present on assayed dendritic segments by the total length of the segments. At least three dendritic regions of interest were analyzed per cell from at least three independent cultures in each experimental condition.

### Live imaging of MT dynamics at spines

Neurons grown on 35 mm glass bottom live imaging dishes (MaTek) were co-transfected with plasmids encoding either EB3-GFP and DsRed or EB3- tdTomato and eGFP using Lipofectamine 2000 (Invitrogen). Live cell imaging was performed 24-48 h after transfection in complete HBSS media (HBSS, 30 mM glucose, 1 mM CaCl2, 1 mM MgSO4, 4 mM NaHCO3, and 2.5 mM HEPES, pH 7.4) using an IX83 Andor Revolution XD Spinning Disk Confocal System. The microscope was equipped with a 100×/1.49 oil UApo objective, a multi-axis stage controller (ASI MS-2000), and a controlled temperature and CO2 incubator. Movies were acquired with an Andor iXon Ultra EMCCD camera and Andor iQ 3.6.2 live cell imaging software. Movies of MT dynamics at spines were acquired at 4 s/frame for 10 min with 3 z-stack planes at 0.4 um step size. Maximum projections of movies were performed by Image Math within Andor software, exported as Tiff files and analyzed in ImageJ. Kymographs were generated by drawing a region from the base of the spine to the tip of spine head. Parameters describing MT invading into spines were defined as follows: % of spines invaded 10 min^-1^: number of spines invaded by MTs during 10 min movie/total number of spines in the imaging field; invasion lifetime: total duration of EB3 residing in a spine including comet lifetimes of multiple invasions ^10^. Parameters describing MT dynamics were defined as follows: rescue/nucleation frequency: number of rescue or nucleation events per μm^2^ per min; catastrophe frequency: number of full tracks/total duration of growth; comet density: number of comets per μm^2^ per min; growth length: comet movement length in μm; comet lifetime: duration of growth; growth rate: growth length/comet lifetime ^91^.

### Analysis of spine structural plasticity

Morphologies (stubby, mushroom, thin) of all protrusions invaded or not invaded by EB3 in the same imaging field before (0 h) and after vehicle or oAβ treatment (2 h) were individually documented using NeuronStudio Software. Percentages of the same protrusions changing to pruned, thin, mushroom, or stubby spines were then calculated based on total number of spines invaded or not invaded by EB3 in the same field. χ^2^ tests were performed on spine persistence or pruning in vehicle and oAβ treated neurons at 0 and 2 hours. χ^2^ tests were also performed on spine morphology changes (to thin, to stubby, to mushroom, to pruned) in vehicle and oAβ treated neurons at 0 and 2 hours.

### Biochemical analysis of post-mortem human brain tissues

Human brains were provided by the Human Brain Tissue Bank, Semmelweis University, Budapest, Hungary. Tissue samples consist of 4 regions of brain (entorhinal cortex, hippocampus, temporal and lateral prefrontal cortex) coming from a panel of 29 male and female patients aged from 52 to 93 years: 11 controls, 5, 6 and 7 from each group corresponding to Braak stadium I-II, III-IV and IV-V (Table S1). <u>Extraction:</u> Brain samples were homogenized 2 x 30 seconds at room temperature in (10% vol / w) 10 mM Tris, 0.32 M sucrose, pH 7.4 containing complete inhibitors cocktail (Roche) using ready to use Precellys Lysing Kit (Bertin Technologies) in a Minilys apparatus. After lysis, the homogenates were collected, frozen in liquid-nitrogen and then stored at −80°C until use. When needed, frozen aliquots were diluted v/v with RIPA buffer (50 mM Tris, 150 mM NaCl, 1% NP40, 0.5% deoxycholate, 0.1% SDS, pH=8) stirred 30 min at 4°C and then centrifuged 10 min at 14000 g at 4°C. Supernatants were frozen in liquid N2 and then stored at −80°C until use. <u>Antibodies:</u> Monoclonal rat anti-tyr-tubulin (YL1/2), polyclonal anti detyrosinated and Δ2 tubulin antibodies were produced in the Andrieux’s lab as previously described ^32^. Mouse monoclonal anti-TTL antibody ID3 was as described ^92^ and polyclonal antibody 13618-1-AP was purchased from Proteintech. <u>Western blot analysis and quantification:</u> RIPA supernatants (10 µl) were subjected to electrophoresis on stain free 4%-15% gels (Bio Rad) and then quickly transferred to Nitrocellulose using Trans-Blot Turbo Transfer System (Bio Rad). Proteins on the membrane were revealed using specific antibodies against different forms of modified tubulin (tyrosinated, detyrosinated, Δ2 tubulin). Anti Tyr-Tub (1/10000), anti deTyr-Tub (1/20000) and anti Δ2-Tub (1/20000) antibodies were used with the appropriated peroxidase-labeled secondary antibodies. Secondary antibody signal was revealed using Pierce ECL Western blotting substrate (Thermo scientific) and analyzed with ChemiDoc™MP Imaging System (Bio Rad) using Image Lab software (stain free gel protocol) for quantification. For each lane of the blot, the software measures the integrated volume of the band corresponding to the antigen of interest. The signal is then normalized according to the total protein measured in the same lane. For every blot, one lane is dedicated to an internal standard corresponding to a WT sample (used for the entire study) and the protein-normalized signal of this standard is considered as 100%, therefore each unknown sample is calculated as a % of this standard. For each brain sample, 3 independent blots were performed and the mean intensity was calculated. <u>ELISA:</u> The assay was routinely performed in high binding 96-wells plates (Immulon 4 HBX, Thermo Fisher). Washings throughout the assay were: 200 μl/well, three times per washing step with PBS buffer solution containing 0.05% Tween 20). Anti-TTL antibody ID3 was coated at 1/2000 in PBS (100 μl/well) overnight (∼ 16 h) at 4 °C. After washing, the plates were blocked by adding 2% BSA in PBS (200 μl/well) for 6 h at room temperature. The plates were then washed and incubated overnight (∼ 16 h) at 4 °C with TTL standards or brain samples diluted in 1% BSA in PBS (100 μl/well). The sample diluent served as negative control. Washed plates were then incubated for 1 h at room temperature with anti-TTL antibody (13618-1- AP) at 1/2000 in 1% bovine serum albumin in PBST (BSA/PBST) (100 μl/well). Washed plates were incubated for 1 h at room temperature with peroxidase rabbit antibody diluted 1:10000 in BSA/PBST (100 μl/well). The plates were washed and incubated with 3,3′,5,5′-tetramethylbenzidine (TMB) Liquid Substrate (Sigma-Aldrich) (100 μl/well). Reaction was stopped after 5 min by adding Stop Reagent for TMB Substrate (Sigma-Aldrich) (100 μl/well). Absorption was determined at 450 nm on Pherastar FS (BMG Labtech). For each brain sample, 3 independent ELISA were performed and the mean value was calculated.

### Biochemical analysis of cultured mouse neurons

Cortical neurons (17 DIV) transduced or not with a lentivirus expressing TTL and treated with DMSO or with 100 nM oAβ (48h) were collected, washed with phosphate-buffered saline medium at 37◦C and directly lysed in Laemmli buffer. The protein contents of TTL, tyrosinated and detyrosinated tubulin were analyzed by quantitative western blot with the protocol, used for human brain samples, described above. Several neuronal cultures were used as indicated in the figure legends and for each sample, 3 independent blots were performed.

### Biochemical analysis of mouse brain tissues

Mice hippocampi were homogenized in a lysis buffer (phosphate buffer saline (PBS) without CaCl2 and MgCl2, 14190-094 Life Technologies) supplemented with protease (P8340, Sigma) and phosphatase inhibitor cocktails (P5726 and P0044, Sigma) at 150mg/mL, using a Precellys apparatus homogenizer (2 x 20s, 5000rpm). Lysates were then centrifuged at 21,000 *g* for 20 min at 4°C. The resulting supernatants were collected and protein concentrations were determined using bicinchoninic acid assays (Pierce/Thermo Fisher Scientific). Samples were stored at −80°C until analysis. <u>Automated western blotting</u> was performed with equal concentrations of protein per sample (0.125 µg/µL) using Peggy Sue™ system (Protein Simple, San Jose, CA, USA) according to the manufacturer’s instructions. Detection of tyrosinated, detyrosinated tubulins and TTL levels were assessed using appropriate primary antibodies as detailed above for human samples. Data were analyzed with Compass software (Protein Simple). Protein levels were normalized using GAPDH signal.

### Immunohistochemical analysis of *post-mortem* brain tissues

De-identified human autopsy brain tissue was obtained from the New York Brain Bank at Columbia University (New York, NY, USA). Neuropathologically-confirmed AD cases and controls were processed following published protocols ^93^. <u>Antibodies</u>: Anti-Δ2 tubulin (AB3203) was from Millipore, anti detyrosinated tubulin (MAB5566) from Sigma Aldrich and anti tau AT8 (MN1020) from Invitrogen. <u>Immunolabelling</u>: brain paraffin blocks were cut into 5 μm sections and deparaffinized in xylene (7 min 2X) followed by 95% EtOH, 90% EtOH, 80% EtOH and 70% EtOH (5 min each). After washing the slices in distilled H_2_O 3X, citric acid was used to retrieve antigen by boiling samples for 15 min. Sections were cooled for 15 min, washed 3X with PBS and blocked with serum for 1 h at room temperature prior to staining with primary antibodies (anti-detyrosinated tubulin, 1/100; anti Δ2tubulin 1/500 and AT8 anti Tau, 1/500) at 4⁰C overnight. The next morning sections were washed 3 X with PBS and stained with appropriate secondary antibodies (Cy3 donkey anti mouse, 1/200; Alexa 488 donkey anti rabbit, 1/200; DAPI, 1/1000) for 1 h at room temperature. Stained samples were washed 3 X with PBS and incubated in 0.1% black Sudan in 70% EtOH for 5 min to reduce auto-fluorescence of lipofuscin, rinsed with 70% EtOH until black was gone and rehydrated in distilled H_2_O. <u>Image acquisition and analyses</u>: Coverslips were mounted with Fluoromount prior to imaging using an Oympus VS-ASW FL 2.7(Build 11032) slide scanner and Olympus soft imaging solutions camera XM10. Images were taken using a 10x objective and same exposure time was used for the same primary antibody (detyrosinated tubulin: 100ms; Δ2: 200ms; AT8 tau: 10ms; DAPI: 10ms). The images were converted into Tiff files for analysis using MetaMorph software. Pyramidal neuron cell bodies and proximal dendrites were randomly selected in the anterior hippocampal formation and average fluorescence intensity was measured for detyrosinated and Δ2 tubulins, as well as for AT8. An average of 150 neurons were selected for each case. Pyramidal neurons were arbitrarily classified into low AT8 (1- 300 A.U.) intermediate AT8 (300.01-1000 A.U.) and high AT8 (1000.01-2400 A.U.) based on AT8 staining intensity in the cell body.

### APPLon and isogenic control iPSC cells maintenance and differentiation

Human induced pluripotent stem cells (hiPSCs) in which the APPV717I (London) mutation was knocked into one allele of the control IMR90 cl.4 iPSC line (WiCell) ^94–96^ using CRISPR/Cas9 was generated by Dr. Andrew Sproul’s lab, as has been described previously ^83^. <u>Maintenance:</u> APPLon knockin (cl. 88) and the isogenic parent line were maintained feeder-free in StemFlex media (Life) and Cultrex substrate (Biotechne). <u>Neuronal differentiation:</u> bankable neural progenitors (NPCs) were first generated using manual rosette selection and maintained on Matrigel (Corning) as has been described previously ^83, 84^. Terminal differentiations were carried out by plating 165,000 - 185,000 NPCs per 12 well plate in N2/B27 media (DMEM/F12 base) supplemented with BDNF (20 ng/ml; Biotechne) and laminin (1 µg/ml; Biotechne) on PEI (0.1%; Sigma) / laminin (20 µg/mL)-coated plates. After 1 week of differentiation, 100 nM AraC (Cytosineβ-D-arabinofuranoside hydrochloride; Sigma) was added to reduce proliferation of remaining NPCs ^84^. A similar strategy was used for imaging plates (MaTek Lifesciences). Differentiations were analyzed 30-40 d post plating. For later passage NPCs, we employed a CD271-/CD133+/CD184+ (Biolegend) FACS purification strategy to remove minority neural crest contaminants (CD271+) that can expand over time, as previously done^97^. <u>Western blot analyses of reprogrammed cortical neurons.</u> Cell lysates from WT and mutant human cortical neurons at 30-40 d of differentiation were lysed in Laemmli sample buffer and boiled at 96°C for 5 min. Cell lysates were sonicated by probe sonication to sheer cellular debris and genomic DNA. Proteins were separated by 10% Bis-Tris gel (Invitrogen) and transferred to nitrocellulose membrane. After blocking in 5% milk/TBS or BSA/TBS, membranes were incubated with primary antibodies (anti total tau (tau 46) (sc-32274) from Santa Cruz; anti tau AT8 (MN1020) and anti-GAPDH (MA5-15738 and PA5-85074) from Invitrogen; anti TTL (13618-1-AP) from Proteintech; anti detyrosinated tubulin (MAB5566) from Sigma Aldrich; Anti-Δ2 antibody (AB3203) from Millipore) at 4°C overnight and 1 h with appropriate secondary antibodies (LI-COR Biosciences). Image acquisition was performed with an Odyssey infrared imaging system (LI-COR Biosciences) and analyzed with Odyssey software.

### Statistical analysis

Data analyses, statistical comparisons, and graphs were generated using GraphPad prism or the R programming language. Statistical analysis between two groups was performed using Student’s t tests for populations with Gaussian distribution or Mann Whitney’s test. Comparison among 3 or more groups was performed using one- or two-way ANOVA with post hoc testing as indicated in text or figure legends (Uncorrected Fisher’s Least Significant Difference, Tukey’s, Dunnett’s or Holm-Sidak’s multiple comparisons test). For comparisons involving two factors and unbalanced samples type 2 or type 3 tests were used. For analysis of data in Fig. 3 B-E, linear mixed models were constructed using the lmer function of the R programming language. Model coefficients including the variance associated with random effects were calculated by restricted maximum likelihood estimation. The distribution of model residuals was verified in graphs of residuals vs. fitted values. The significance of fixed effects was tested in a type II F test of the full model against smaller models (Wald test) using the *Anova* function of the R *car* package. Post-hoc testing of pairwise differences with control was performed by Dunnett’s test on non-weighted marginal means over the relevant factor, using the R *emmeans* package.

Mean differences were considered significant at p < 0.05 (* *p* < 0.05; ** *p* < 0.01; *** *p* < 0.001 and **** *p* < 0.0001). Some exact p values are indicated in text or figures.

