## Supplementary material for "Impaired α-tubulin re-tyrosination leads to synaptic dysfunction and is a feature of Alzheimer’s disease": Supplemtary Information Peris et al

**Peris L. et al.**

### S data

### SUPPLEMENTARY MATERIAL AND METHODS

**Behavioral studies.** Behavioral tests were done in 3 to 4-month old WT and TTL$^{+/-}$ mice. Evaluation of sensorimotor function was performed by analyzing the body weight, water and food intake, muscular tone balance (tail elevation, hanging wire, pole test, chimney test and ring test), and locomotor activity. Evaluation of spatial memory was performed with Morris water maze test. **Body weight**. The mice were weighed with a calibrated scale, at regular intervals. **Locomotor activity**. Spontaneous locomotor activity was measured by using mice individually placed in Plexiglas open-field boxes (L 21.5 x W 12 x H 18 cm) equipped with infrared sensors for accurate location of the animal (Actimeter, IMETRONIC, France). The rack containing 8 boxes was connected to an electronic interface, which provides the formatting of signals from infrared sensors and allows communication with the computer. Each box remained independent and was equipped with 3 parallel infrared photocell units, all located 3 cm above the cage bottom at even intervals along the long axis of the cage, in order to assess movements within the horizontal plane. Ten additional photocell units were placed 10 cm above the cage bottom, at even intervals along the long axis of the cage, to record the frequency of rearing with which the mice stood on their hind legs in the field (vertical activity). Locomotor activity was assessed by counting the number of interruptions of the horizontal photocell units located 3 cm above the cage bottom. This parameter was automatically computed. Spontaneous locomotor activity was recorded in 10 minutes intervals for 60 min. **Rotarod**. The rotarod test is used to assess motor coordination, ataxia and balance in young adult male mice. Mice had to keep their balance on a rotating rod. The time (latency) taken by the mouse to fall down the rod was measured. In the morning of test day, mice were trained to walk on the rotating rod (rotarod apparatus, LETICA, BioSeb) under continuous speed of 40 rpm during 5 min. During the training session, in the afternoon the mice were put back in the rotating rod with an increasing speed (from 4 to 40 rpm). **Water and food intake.** The mice were placed in a cage, with the same stall as the supplier. The food and water were weighed before the animals are brought together. After 24 h or 3 days, the food and water were weighed again. **Muscular tone balance**. Tail elevation test: the mice was suspended by the tail and the spontaneous position of the limbs was observed during 5 s. The normal position is with limbs in an escaped extended position with no clasping whereas abnormal position is with limbs entirely retracted and touching the abdomen (scores between 5 and 0). Wire hang test: the mice were placed in the center of the sieve about 10 cm above a table. The sieve is slowly turned over and the falling latency is recorded during 3 tests lasting a maximum of 2 min with 10-15 s intervals. Pole test: the mice were placed at the top of a vertical wooden pole (height 60 cm) and the ability of the mice to grasp, to descend to the base of the pole was evaluated in 3 consecutive trials. Behavior is ranked normal when the mice perfectly climbing down with 4 paws, intermediate when the mice is climbing down with difficulties and abnormal when the mice do not grip the test pole (scores between 5 and 0). Chimney test: the mice were introduced head forward in a pyrex glass cylinder (3 cm diameter and 30 cm length) held in a horizontal position. The tube is then moved to a vertical position and the ability of the mice to climb backwards out was monitored. Behavior is ranked normal when the mice use the 4 legs and reach the top of the tube quickly, intermediate when the mice reach the top of the tube with difficulty and abnormal when the mice cannot go backwards (scores between 5 and 0). Ring test: the mice were put on a metal ring and the locomotor behavior was monitored. Behavior is ranked normal when the mice perfectly climb and grip the ring with 4 paws, intermediate when the mice grip with difficulties and abnormal when the mice do not grip the ring (scores between 5 and 0).

**Morris water maze test** was conducted in a pool filled with white tinted water that contain a submerged platform always placed at the same place 1 cm below the surface of the water. During learning, mice are subjected to several tests in the pool, in which they must learn, starting from different locations (N, S, E and W), to swim to the platform guided by marks available outside the pool. After 24 h, the mice are subjected to a retention test to evaluate their spatial memory. The percentage of time spent in the goal quadrant was measured for each animal the mean and standard errors of the mean (SEM) were calculated for each group.

### SUPPLEMENTAL FIGURES

#### Figure S1

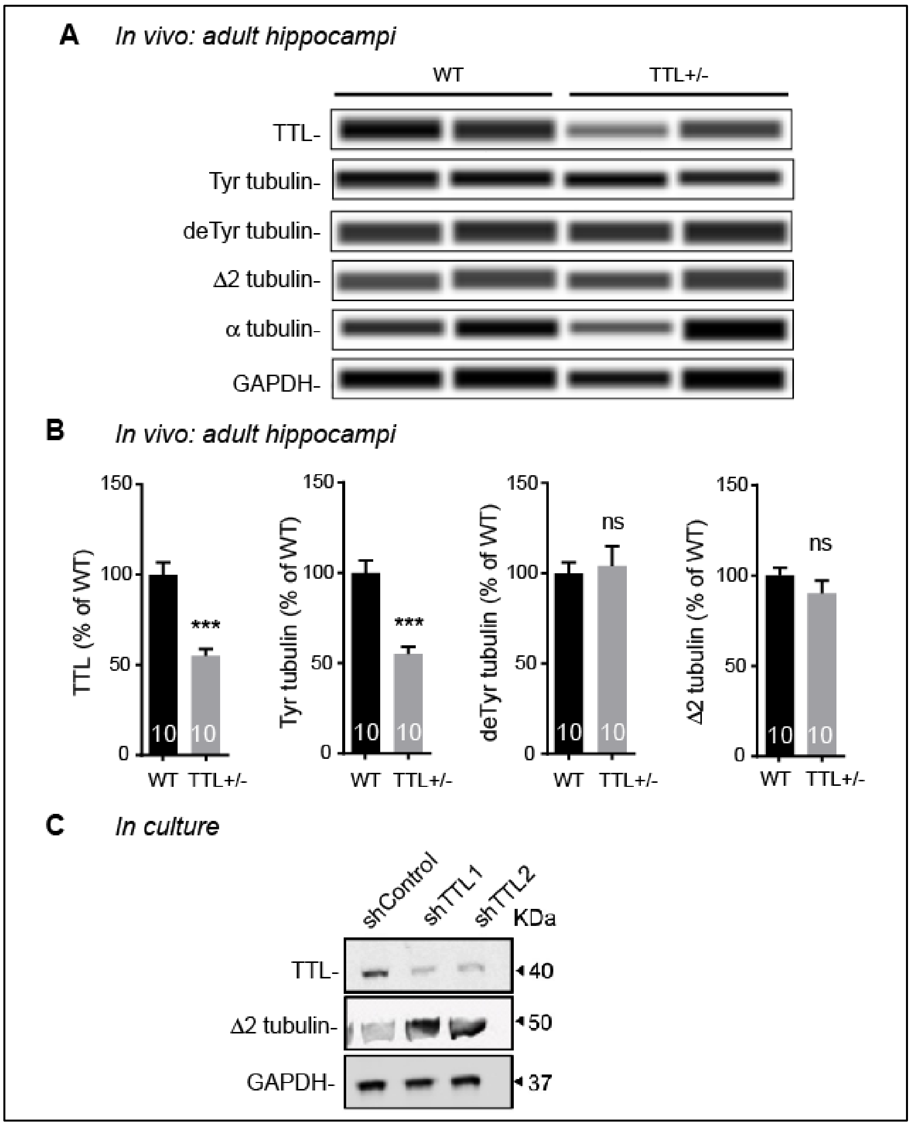


**Figure S1. Analysis of TTL levels and α-tubulin modifications linked to the Tyr/deTyr cycle in protein extracts from WT and TTL$^{+/-}$ hippocampi and in WT neurons silenced of TTL expression.**

(**A-B**) PeggySue analysis of TTL and selected modified tubulin levels in 3-month-old WT and TTL$^{+/-}$ mouse hippocampi (samples preparation and analysis as in main text, Fig 1B). (A) Examples of protein expression levels. (B) Quantification of relative amounts of proteins. All values are normalized with GAPDH and expressed as % of WT. Graphs represent mean ± SEM. n = number of animals analyzed are shown in each graph. Mann-Whitney's test *** $p < 0.001$ (**C**) Immunoblot analysis of TTL, Δ2 tubulin and GAPDH levels in cultured rat hippocampal neurons infected with control shRNA (non-coding) or 2 independent shRNA lentiviruses targeting TTL (shTTL1 and shTTL2).

**Figure S2**

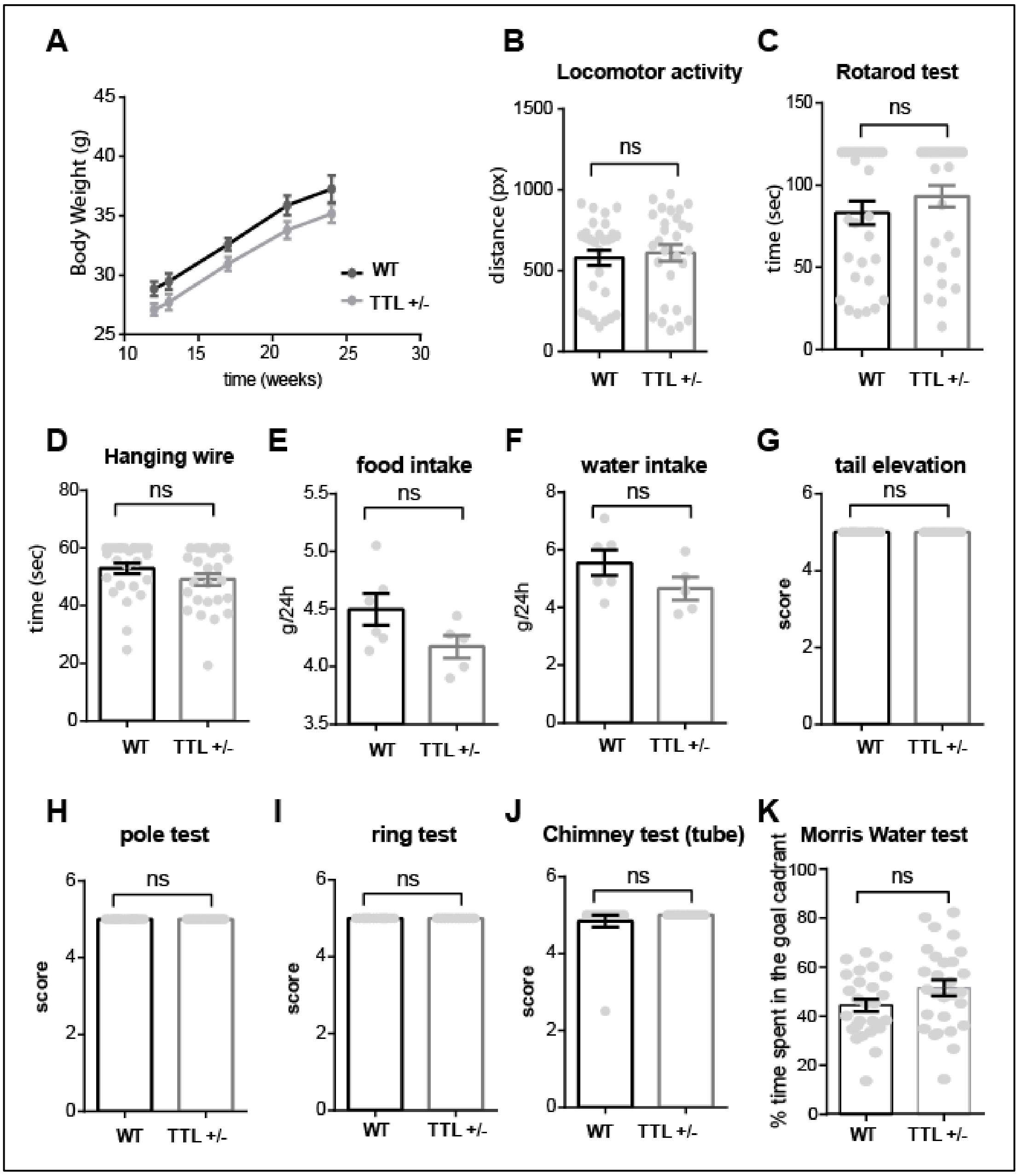


**Figure S2: Normal sensorimotor function, locomotor activity and spatial memory of $TTL^{+/-}$ mice.** (**A**) Body weight of WT and $TTL^{+/-}$ mice was measured in function of time. Two Way ANOVA, genotype x time interaction ($F (4, 88) = 0.0681$, $p = 0.9914$). n = 12 WT and $TTL^{+/-}$ mice. Source of variation: genotype, * $p = 0.020$. (**B**) Spontaneous locomotor activity was analyzed by measuring total distance covered by WT and $TTL^{+/-}$ mice in open field boxes in 30 minutes. n = 28 WT and $TTL^{+/-}$ mice. Student t test, ns = not significant. (**C**) Motor coordination was analyzed by rotarod test. The time taken by the mouse to fall down the rod was measured for WT and $TTL^{+/-}$ mice. n = 30 WT and $TTL^{+/-}$ mice. Student t test, ns = not significant (**D-E**). Water and food intake were measured by weighing water and food at the beginning of the test and 24 h later, for WT and TTL +/- mice. n = 6 WT and 5 $TTL^{+/-}$ mice.

Student t test, ns = not significant. **(F-J)** Fine motor coordination was analyzed by various tests: (F) Hanging wire test (n = 26 WT and $TTL^{+/-}$ mice, Student t test, ns = not significant), (G) Tail elevation, (H) Pole test, (I) Ring test and (J) Chimney test (n=16 WT and 15 $TTL^{+/-}$ mice, ns = not significant). (**K**) Evaluation of spatial memory was performed with Morris water maze test. The percentage of time spent in the goal quadrant was measured for WT and $TTL^{+/-}$ mice. n = 27 and 28 for WT and $TTL^{+/-}$ mice. Mann-Whitney test, ns = not significant. (**B-K**) All graph represents mean ± SEM.

**Figure S3**

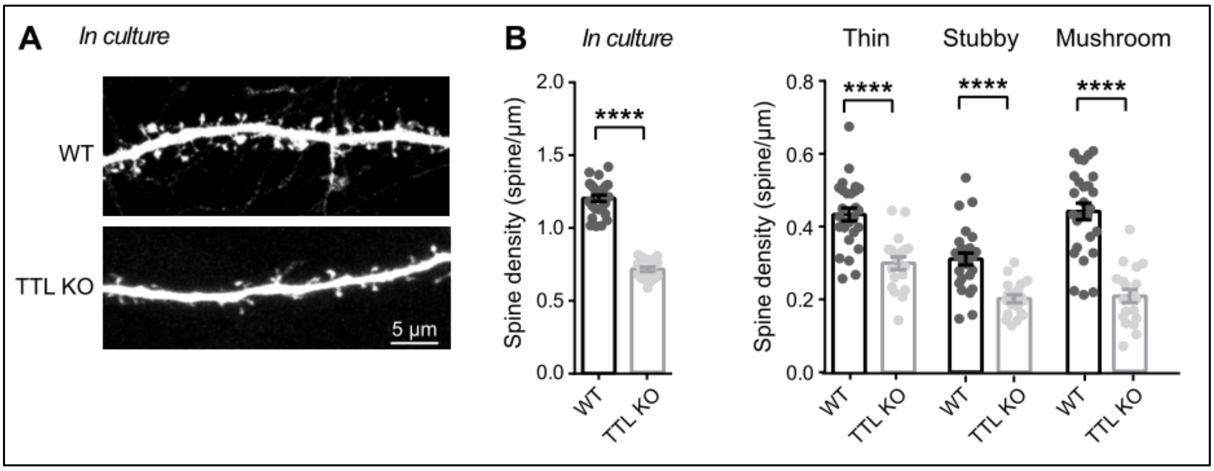


**Figure S3. Total TTL suppression results in dramatic loss of dendritic spine density*.***

**(A)** Confocal images showing representative examples of dendritic segments of GFP-expressing WT and TTL KO hippocampal neurons in culture at 17 DIV. **(B)** Total dendritic spine density, or that of each different morphological type of spines are represented from WT and TTL KO hippocampal cultured neurons. Graphs represent mean ± SEM. n = 27 and 19 neurons from WT and TTL KO embryos from at least 3 independent cultures. Student's t test. **** $p$<0.0001.

**Figure S4**

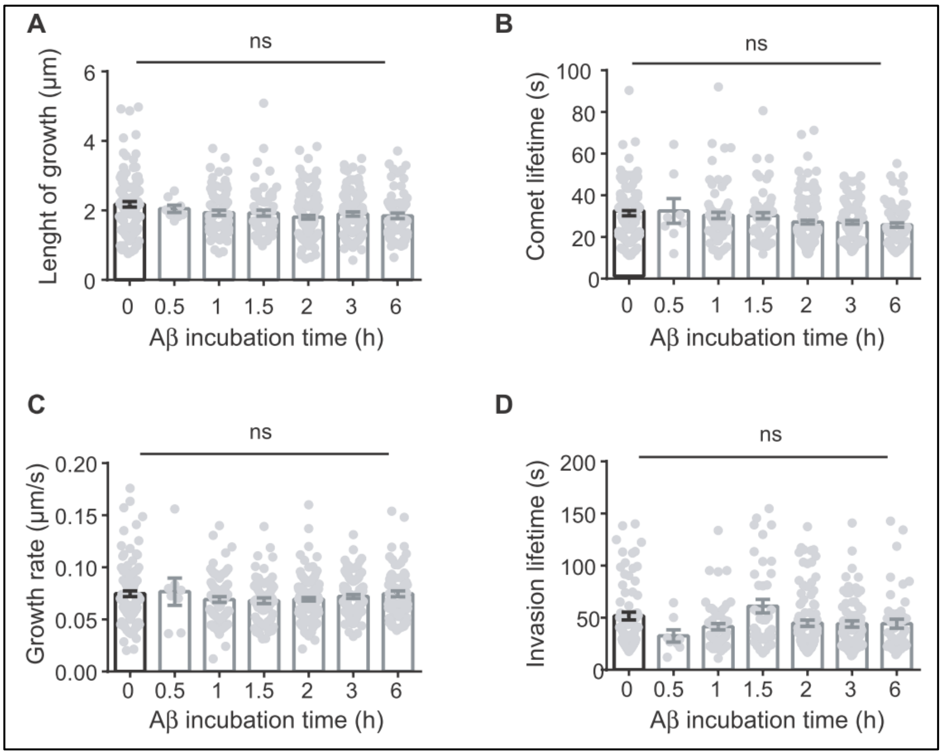


**Figure S4. Dynamic parameters of MT invading spines before and after oAβ treatment**. Comet length growth (A), comet lifetime (B), MT growth rate (C) and invasion lifetime (D) during 10 min movies in WT hippocampal neurons expressing EB3-EGFP and DsRed before and after oAβ (250 nM) incubation at the indicated times. Graphs represent mean ± SEM. n=112, 8, 76, 66, 138, 104, and 73 comets were analyzed at each time point, respectively. Kruskal-Wallis test, ns= not significant.

**Figure S5**

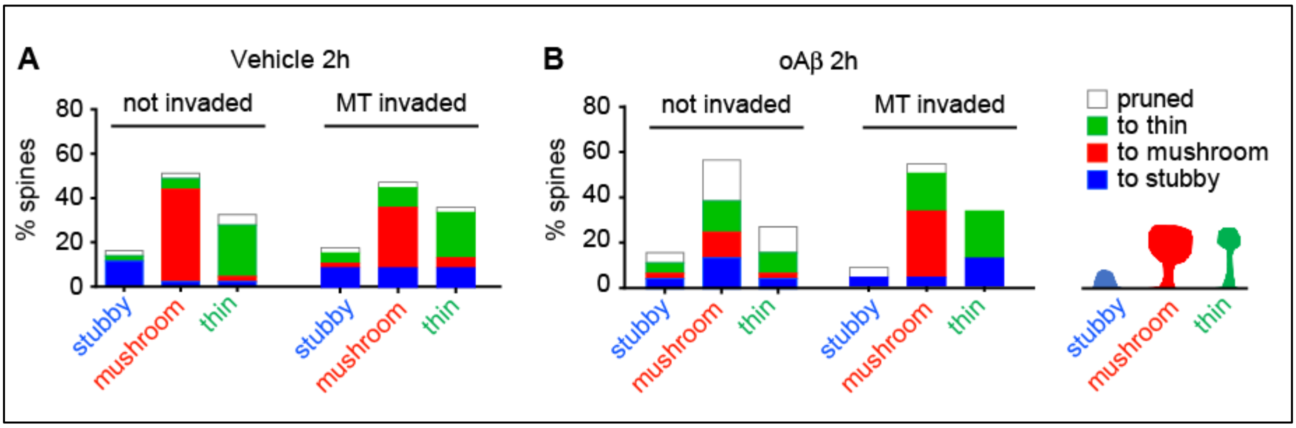


**Figure S5. Structural plasticity of spines invaded or not by dynamic MTs.**
Morphologies (stubby, mushroom, thin) of all dendritic protrusions invaded or not invaded by EB3-labeled growing MT plus ends before and after vehicle (A) or 250 nM oAβ treatment (B) were individually documented. Percentages of the same protrusions changing to pruned, thin, mushroom, or stubby spines were then calculated based on total number of spines invaded or not invaded by EB3 in the same field. The $\chi^2$ test was performed on the 4 conditions (Veh 2 h not invaded, Veh 2 h MT invaded, oAβ 2 h not invaded and oAβ 2 h MT invaded) and the 4 possible morphological outcomes (to stubby, to mushroom, to thin, to pruned). The fates of the spines between conditions were significantly different ($\chi^2$ = 53.98, 9 df, p = < .0001).

### Supplemental Tables

#### Table S1

| Control patients | | | | |
|---|---|---|---|---|
| Patient Number | | Gender | Age | Cause of death |
| #81 | Control | male | 65 | pulmonary embolism, arterosclerosis, heart failure |
| #199 | Control | female | 76 | heart failure |
| #223 | Control | male | 61 | heart failure |
| #220 | Control | male | 63 | pulmonary embolism |
| #164 | Control | male | 85 | acute cardiorespiratoric insufficiency |
| #198 | Control | female | 68 | parieto-occipital and frontal stroke, pneumonia, respiratory insufficiency |
| #205 | Control | male | 66 | cardiovascular-pulmonary insufficiency, acute lymphoid leukemia |
| #217 | Control | male | 60 | acute myocardial infarction |
| #266 | Control | male | 61 | vertebrobasilar stroke (right side) |
| #269 | Control | male | 79 | stroke in the left cortical hemisphere, herniatio |
| #276 | Control | female | 79 | stroke, pneumonia |
| **Alzheimer's disease patients** | | | | |
| #184 | Braak stadium I - II | male | 83 | emolitio cerebri |
| #187 | Braak stadium I - II | male | 62 | respiratory and cardiac insufficiency |
| #191 | Braak stadium I - II | female | 93 | Alzheimer's disease, cardiovascular-respiratory insufficiency |
| #218 | Braak stadium I - II | female | 72 | acute cardiac insufficiency |
| #219 | Braak stadium I - II | male | 67 | pulmonary embolism |
| #185 | Braak stadium III - IV | male | 80 | stroke (right side), herniatio |
| #196 | Braak stadium III - IV | female | 78 | stroke, arteria cerebri media, brain hemorrhage |
| #230 | Braak stadium III - IV | female | 79 | pulmonary embolism |
| #197 | Braak stadium III - IV | male | 64 | myocardial infarction |
| #267 | Braak stadium III - IV | female | 91 | stroke, arteria cerebri media (left side) |
| #279 | Braak stadium III - IV | male | 79 | stroke (right side) |
| #154 | Braak stadium V-VI | female | 72 | acute myocardial infarction, earlier heart failure, arterosclerosis |
| #167 | Braak stadium V-VI | female | 65 | suicide (hanging - asphyxia) |
| #195 | Braak stadium V-VI | male | 83 | respiratory and cardiac insufficiency |
| #202 | Braak stadium V-VI | male | 84 | cardiac and respiratory insufficiency |
| #212 | Braak stadium V-VI | female | 87 | dementia, myocardial insufficiency |
| #229 | Braak stadium V-VI | female | 78 | Alzheimer's disease |
| #232 | Braak stadium V-VI | female | 77 | cardiorespiratory insufficiency |

**Table S1. Details of control and Alzheimer's disease patients classified into Braak stages as shown in Figure 3A-E.** The gender, age and cause of death are described for each patient. As the post-mortem interval (PMI) between the time of death and the collection of tissues is a critical factor affecting the quality of human brain tissues, we analyzed only samples with a PMI inferior of 300 min.

**Table S2**

| Post# | MPN08-17 | MPN03-166 | MPN09-270 | MPN07-47 | OC04-18 | MPN03-161 |
|---|---|---|---|---|---|---|
| **Case & Classification** | Case 2 Control | Case 3 Control | Case 4 Control | Case 9 AD | Case10 AD | Case 11 AD |
| **Age** | 74 | 87 | 90 | 75 | 82 | 86 |
| **Sex** | M | M | M | M | M | M |
| **Braak NFT stage** | 3 | 3 | 3 | 6 | 6 | 6 |
| **CERAD plaque score** | none | sparse | sparse | frequent | frequent | frequent |
| **Amyloid angiopathy** | 0 | 0 | 0 | ++ | ++ | ++ |
| **NIAR** | 0 | Low | Low | High | High | High |
| **PMI (min)** | 93 | 310 | 270 | 295 | 127 | 260 |

**Table S2. Table of case descriptions of postmortem human AD and control brains shown in Figure 3F-H.** The age and gender are listed for each patient. The Braak NFT stage, CERAD plaque score and amyloid angiopathy are listed to demonstrate the pathological hallmarks of AD present in each brain sample. The NIA-Reagan score (NIAR) is a post-mortem diagnosis score of the likelihood of having AD, which takes into account the Braak stage and CERAD score. We analyzed only samples with a PMI less than or close to 310 min.
